# CASTLE: a training-free foundation-model pipeline for cross-species behavioral classification

**DOI:** 10.1101/2025.08.22.671685

**Authors:** Yu-Shun Liu, Han-Yuan Yeh, Yu-Ting Hu, Bing-Shiuan Wu, Yi-Fang Chen, Jia-Bin Yang, Sureka Jasmin, Ching-Lung Hsu, Suewei Lin, Chun-Hao Chen, Yu-Wei Wu

## Abstract

Accurately and efficiently quantifying animal behavior at scale without intensive manual labeling is a long-standing challenge for neuroscience and ethology. Keypoint-based tracking emphasizes simplicity and efficiency but loses the richness of posture and context, while emerging foundation models capture pixel-level details, yet often require nontrivial efforts of retraining and can be more sensitive to backgrounds or lighting. Here, we present *CASTLE*, a training-free pipeline that addresses all these issues by synergistically combining foundation models for segmentation, tracking, and feature extraction. By isolating regions-of-interest (ROI), *CASTLE* first generates "focused (ROI-masked)” and orientation-invariant latent features, capturing rich postural details in zero-shot, fine-tuning-free manners. Following ROI isolation, *CASTLE,* through an interactive “Behavior Microscope” module, supports hierarchical clustering, for progressive, human-in-the-loop embedding and clustering. This enables raw-image-assisted discovery of behavioral classes without predefined categories. Across mice, *Drosophila* and *C. elegans*, *CASTLE* matches expert class annotations (>90%), reveals disease-relevant phenotypes in Parkinsonian mouse models. By eliminating purpose-specific model training and providing a raw-image-informed accessible workflow, *CASTLE* offers a scalable framework for interpretable, cross-species behavioral phenotyping.

## Introduction

Quantifying behavior with the same rigor that we quantify neural activity is now a central mandate in systems neuroscience^1–3^, psychiatry^4^ and drug discovery^5^. The basic workflow contains image feature extraction (segmentation), tracking and classification, with efficiency, feature complexity, and interpretability the major dimensions of consideration. Traditional analyses rely on manual observation, a labor-intensive process prone to subjectivity and inter-observer variability^5–8^. While deep-learning tools like *DeepLabCut (DLC)*^9^ *and Lightning Pose*^10^ have made keypoint-based pose estimation a practical alternative, they introduce new bottlenecks. Achieving detailed analysis—such as resolving digits or subtle paw rotations—requires extensive labeling and model retraining for each new experiment. Fundamentally, these methods reduce an animal’s complex posture to a sparse skeleton, which can cause unanticipated or fine-grained behavioral patterns to be overlooked.

To move beyond skeletonized sparse representations, recent approaches apply Visual Foundation Models (VFMs)^11^ directly to raw videos, using tools like *Animal-JEPA*^12^, *VideoPrism*^13^ and *MouseGPT*^14^ to learn dense embeddings that preserve rich postural details. Yet, these powerful VFMs present a new dilemma. Most VFM workflows still require task-specific fine-tuning or a supervised classifier to link embeddings to behavior. More importantly, their broad representations of image details often work against behavioral analysis with specific purposes; unless explicitly focused, VFMs encode background clutters, lighting changes, orientation, etc., adding nuisance variance that masks the signals of interest. A final hurdle is interpretation: with often used unsupervised algorithms, the complex, high-dimensional data from VFMs can be hard to understand, yielding classifications that are mathematically plausible but lack biological interpretability. This underscores the need for an integrated pipeline that extracts focused features, and provides a robust framework for interpreting them.

To balance efficiency, feature complexity and behavioral interpretability, we introduce *CASTLE* (*C*ombined *A*pproach for *S*egmentation and *T*racking with *L*atent *E*xtraction and exploration), a pipeline designed to leverage different foundation models for their complementary roles. *CASTLE* integrates zero-shot segmentation (*Segment Anything Model* (*SAM*)^15^), robust tracking (*DeAOT*^16^), and dense visual feature extraction (*DINOv2*^17^). After segmenting the image for each region-of-interest (ROI), *CASTLE* applies an ROI to produce "focused" latent features that ignore background noise and are invariant to animal orientation. As the result, this pipeline is training-free, requiring no task-specific retraining or fine-tuning while preserving complex image features. To facilitate interpretation, *CASTLE* includes a powerful GUI-assisted interface called “Behavior Microscope” for hierarchical clustering of features. This tool empowers researchers to directly visualize the latent feature space, examine video segments corresponding to each cluster, and iteratively refine classification. This human-in-the-loop validation ensure that the resulting behavioral categories are not only computationally robust but also biologically comprehensible, bridging the gap between high-dimensional image data and ethological insight. Thus, human decisions are involved only close to the most semantic level—toward classification—which realizes balanced efficiency and interpretability. Crucially, however, human intuition is helping, not limiting, behavioral pattern discovery through hierarchically ‘peeling off’ well recognized data and leaving the more challenging, less understood clusters to be identified. Objectivity, decision process and flexibility complement one another in our design principle.

We validated *CASTLE* across diverse contexts—including skilled reach-and-grasp in mice, grooming in *Drosophila*, and foraging in *C. elegans*, without intensive manual labeling or network retraining. *CASTLE* has achieved over 90% agreement with expert annotations and uncovered behavioral classes that are not predefined. By managing the intertwined challenges of data visualization, feature extraction, and interpretation, *CASTLE* establishes a scalable and accessible paradigm for behavioral neuroscience.

## Results

### *CASTLE* accurately extracts and identifies discrete behavioral classes from video data

We developed the *CASTLE* pipeline to facilitate precise behavioral classification directly from raw video data (**Fig. 1**). The flowchart of the *CASTLE* pipeline is provided in **Fig. 1a** with detailed mathematic descriptions in **Extended Data Fig. 1a**. Initially, *CASTLE* uses a user-define and semi-automatic segmentation approach where users define regions-of-interest (ROIs) using *SAM*^15^ (**Fig. 1b and Extended Data Fig. 1b,c**), typically corresponding to key anatomical landmarks, such as paws and nose in mice (**Fig. 1c and Extended Data Fig. 1b**). These ROIs are tracked frame-by-frame using the *DeAOT* video object segmentation model^16^, yielding robust spatial tracking that captures movement trajectories and changes in ROI area over time.

**Figure 1.**
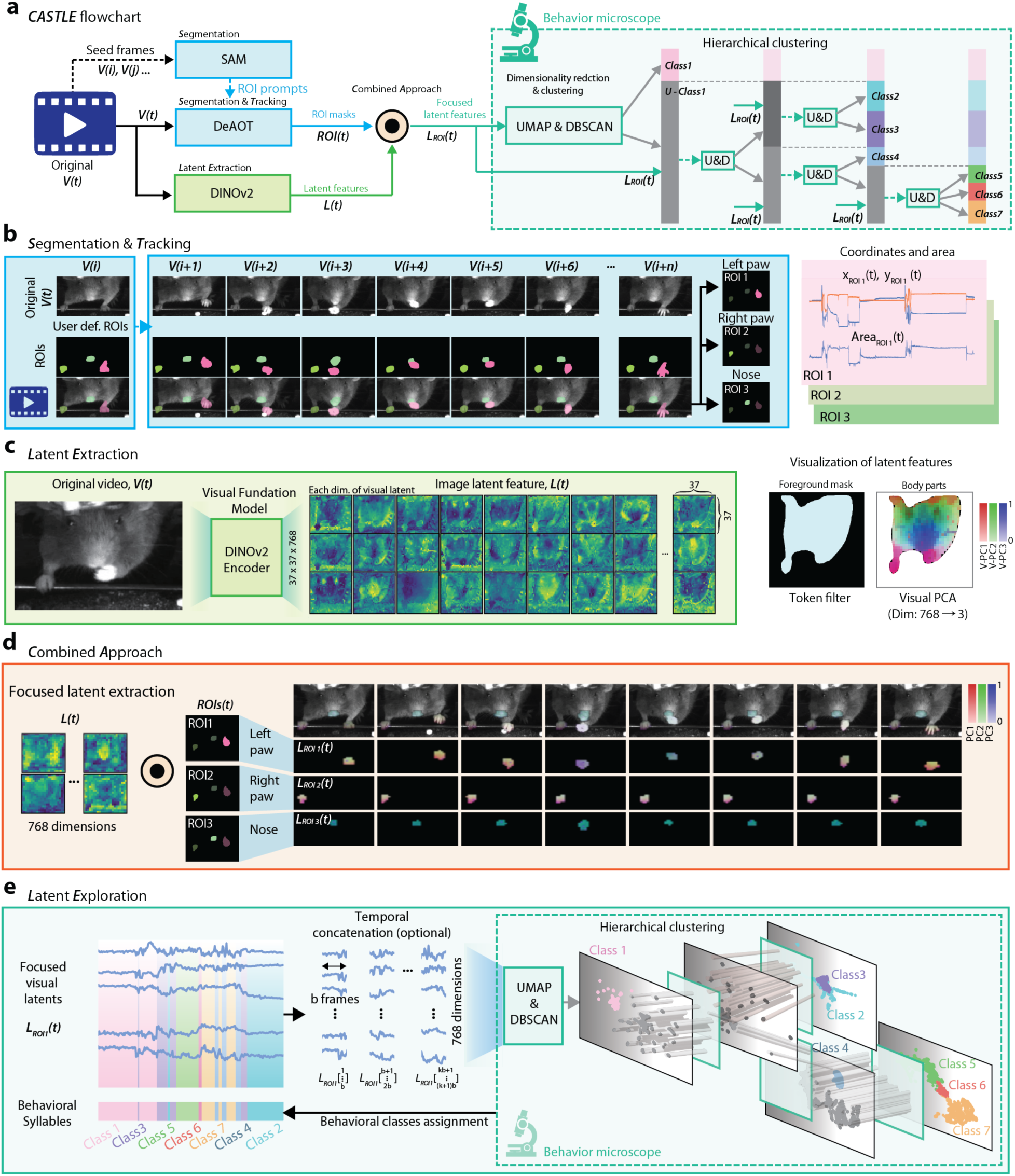
Overview of the *CASTLE* pipeline for deriving behavioral classes from video data. **a. Overview of *CASTLE*’s flowchart.** The pipeline comprises a Combined Approach for Segmentation & Tracking, Latent Feature Extraction, for focused feature generation, and Latent Exploration facilitated by the “Behavior Microscope” for hierarchical clustering and behavioral class assignment. **b**. **Segmentation & Tracking**. The process begins by extracting a selected frame (*V(i)*) of the video with *SAM*, allowing researchers to define specific Regions-of-Interest (ROIs). A video object segmentation model, *DeAOT*, then tracks these ROIs across subsequent frames, providing detailed information on their position and area over time. **c**. **Latent Feature Extraction**. A visual foundation model, *DINOv2*, is employed to encode each video frame into high-dimensional features, capturing comprehensive visual information for the entire image. These features are then integrated with the ROI data from **b** to produce a focused visual latent representation (*L_ROI_(t)*), highlighting key aspects of the tracked objects. **d. Combined Approach**: The focused visual latent features derived from ROIs are used for further behavioral classification. For visualization, the ROI-specific visual latent features were projected onto principal component analysis (PCA) space, generating dimension-reduced RGB images corresponding to the PC1-3 of the visual latent features. Noted that only the moving left paw exhibits clear changes in RGB colors over time. **e. Latent Exploration**. The focused latent representations (*L_ROI_(t)*) are analyzed by comparing similarities between pairs of frames using embedding (*UMAP*) and clustering (*DBSCAN*) techniques. Similar frames are grouped into clusters, which are then assigned to initial behavioral classes. This entire process, including subsequent hierarchical clustering to identify broader behavioral classes along the timeline, is interactively performed and refined using the “Behavior Microscope” interface.

To ensure comprehensive visual semantic capture, *CASTLE* employs *DINOv2*, a state-of-the-art visual foundation model^17^, encoding each video frame into rich 768-dimensional visual latent features (**Fig. 1c**). For visualization and enhancing interpretability, the visual latent features undergo dimensionality reduction via Principal Component Analysis (PCA) reveals a stable representation of body parts of the mouse over different image frames (**Fig. 1c**, *right panel;* **Extended Data Fig. 1d** and **Supplementary Video 1**). These latent representations are combined with ROI-specific spatial data, resulting in "focused visual latent features” that accurately represent the visual dynamics associated with each anatomical feature of interest (**Fig. 1d** and **Supplementary Video 2**). The temporal dynamics of “focused visual latent features” were visualized by PCA colormap (**Fig. 1d**). Notably, only the left paw (ROI1) showed drastic color change over time, indicating the variation in the posture of the left paw but not the right paw (ROI2) and the nose (ROI3). Without focused latent extraction, the first 3 PC features do not reflect clear changes in all body parts (**Extended Data Fig. 1d**), suggesting the importance of the application of “Focusing”.

Finally, *CASTLE* performs latent feature exploration first with an optional step of temporally concatenating the extracted ROI-specific latent signals. Then through *CASTLE*’s “Behavior Microscope” module, the latent signals are embedded into a lower-dimensional space using Uniform Manifold Approximation and Projection (*UMAP*)^18^. Clusters formed through Density-Based Spatial Clustering of Applications with Noise (*DBSCAN*)^19^ define initial behavioral classes, followed by human-in-the-loop hierarchical clustering to define behavior classes (**Fig. 1a,e**), which are assigned to the temporal axis to reveal consistent and repetitive behavioral patterns into ethogram (**Fig. 1e**). This comprehensive and efficient pipeline (**Fig. 1a**) thus enables high-throughput behavioral quantification from unannotated video data.

### Intuitive Graphical User Interface (GUI) for behavioral analysis using *CASTLE*

The *CASTLE* pipeline is integrated into a user-friendly GUI, making behavioral analysis streamlined and intuitive. The interface simplifies complex analytical procedures into clearly defined steps. Initially, users employ the segmentation and tracking module (**Fig. 2a, d**) to interactively define and adjust ROIs across video frames. In this module, user input establishes initial tracking conditions, while automated mask predictions facilitate iterative adjustments, ensuring robust and accurate ROI tracking.

**Figure 2.**
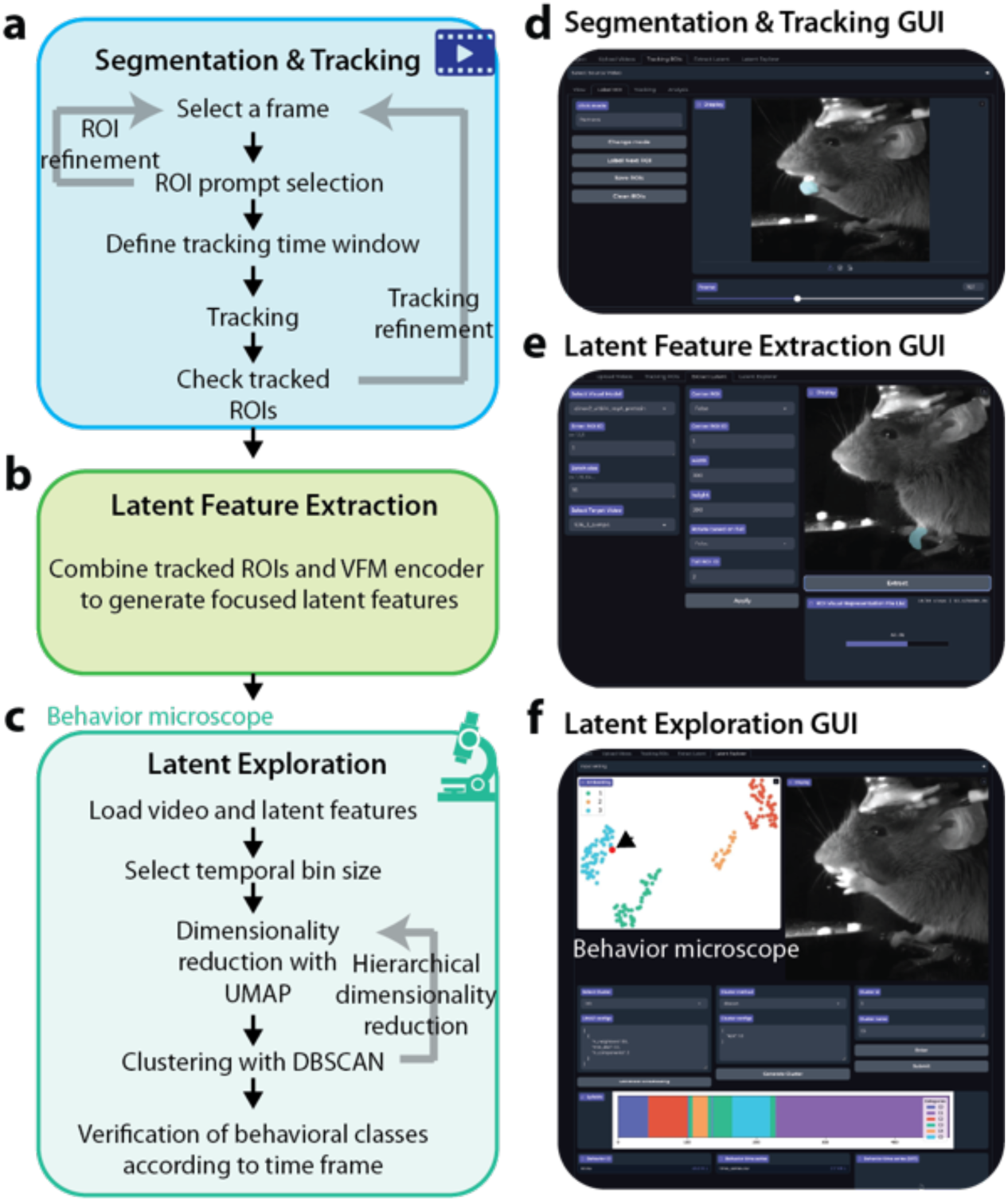
Implementation of the *CASTLE* pipeline within a graphical user interface (GUI) for behavior analysis. **a-c.** Workflow overview in the GUI, showing the three main stages of the *CASTLE* pipeline: **a.** Segmentation & Tracking: Users select a reference frame and define regions-of-interest (ROIs) via interactive prompts. The GUI facilitates the definition of tracking time windows and subsequent refinement of ROI tracking across video frames. **b.** Latent Extraction: This stage integrates video data with tracked ROI information to generate ROI-specific visual latent features, capturing crucial visual aspects necessary for behavioral analysis. **c.** Latent Explorer: Users load the combined latent and video data, select suitable temporal bins, and perform dimensionality reduction (*UMAP*), optionally with hierarchical refinement. Behavioral classification is executed through clustering (*DBSCAN*), enabling temporal verification of classes and identification of behavioral classes. **d-f.** GUI Screenshots: **d.** Segmentation **&** Tracking GUI: Interface for defining, tracking, and refining ROIs. **e.** Latent Extraction GUI: Visualization of how video frames and ROI data combine to form latent features. **f.** Latent Exploration GUI (“Behavior Microscope”): Displays results of dimensionality reduction, cluster analyses, and timelines of behavioral classifications (also see **Extended Data Fig. 2** and **Supplementary Video 3**).

In the subsequent latent feature extraction step (**Fig. 2b, e**), visual data from the video is combined seamlessly with ROI information to generate ROI-specific latent features. These visual latent features effectively capture essential dynamics critical for distinguishing behaviors, forming a powerful intermediate representation for detailed analysis. Lastly, the latent exploration module (**Fig. 2c, f**) offers extensive tools for visualizing and analyzing latent features. Within this module, the “Behavior Microscope” serves as the primary interface for discovery and refinement. Users can load videos with latent data, conduct dimensionality reduction using *UMAP*, and then interactively probe the resulting clusters (**Extended Data Fig. 2 and 3**). This interface is central to the human-in-the-loop workflow, allowing for the recursive refinement of behavioral classes and ensuring that the final ethogram is both data-driven and validated by expert intuition. Additionally, the interface supports recursive refinement through hierarchical clustering, which clearly differentiates behavioral classes, and interactive verification tools allow users to precisely localize and validate behavioral classes (**Extended Data Fig. 2 and 3**). Collectively, the intuitive GUI implementation of *CASTLE* significantly reduces the complexity associated with advanced behavioral analyses, enabling sophisticated latent-space exploration accessible even to researchers without extensive computational expertise.

### Comparison of *CASTLE* and *DeepLabCut* in paw tracking

We aimed to evaluate whether *CASTLE* could achieve tracking performance comparable to established marker-based methods, such as *DeepLabCut* (*DLC*), while providing additional advantages in efficiency and robustness. Comparative analyses between *CASTLE* and *DLC* for paw tracking during mouse reaching tasks revealed similar tracking accuracy. Visual inspection demonstrated comparable trajectories generated by *CASTLE* and those labeled by *DLC*, particularly during significant movements exceeding defined displacement thresholds (**Fig. 3a**). However, trajectories from *DLC* consistently exhibited greater pre-movement displacement and higher noise levels, especially evident during stationary/resting phases across multiple trials (**Fig. 3b**). Such jitter in *DLC*-generated trajectories likely arises from minor variations inherent in manually labeled ground truth data used to train the *DLC* model. Quantitative assessments further supported *CASTLE*’s advantage in noise reduction, showing significantly lower standard deviation values of paw speed during resting states compared to *DLC* (Wilcoxon signed-rank test, p < 0.0001; **Fig. 3c-d**). This reduced jitter underscores *CASTLE*’s robustness and highlights its efficiency, as it eliminates the need for extensive manual labeling or prolonged neural network training periods required by *DLC*.

**Figure 3.**
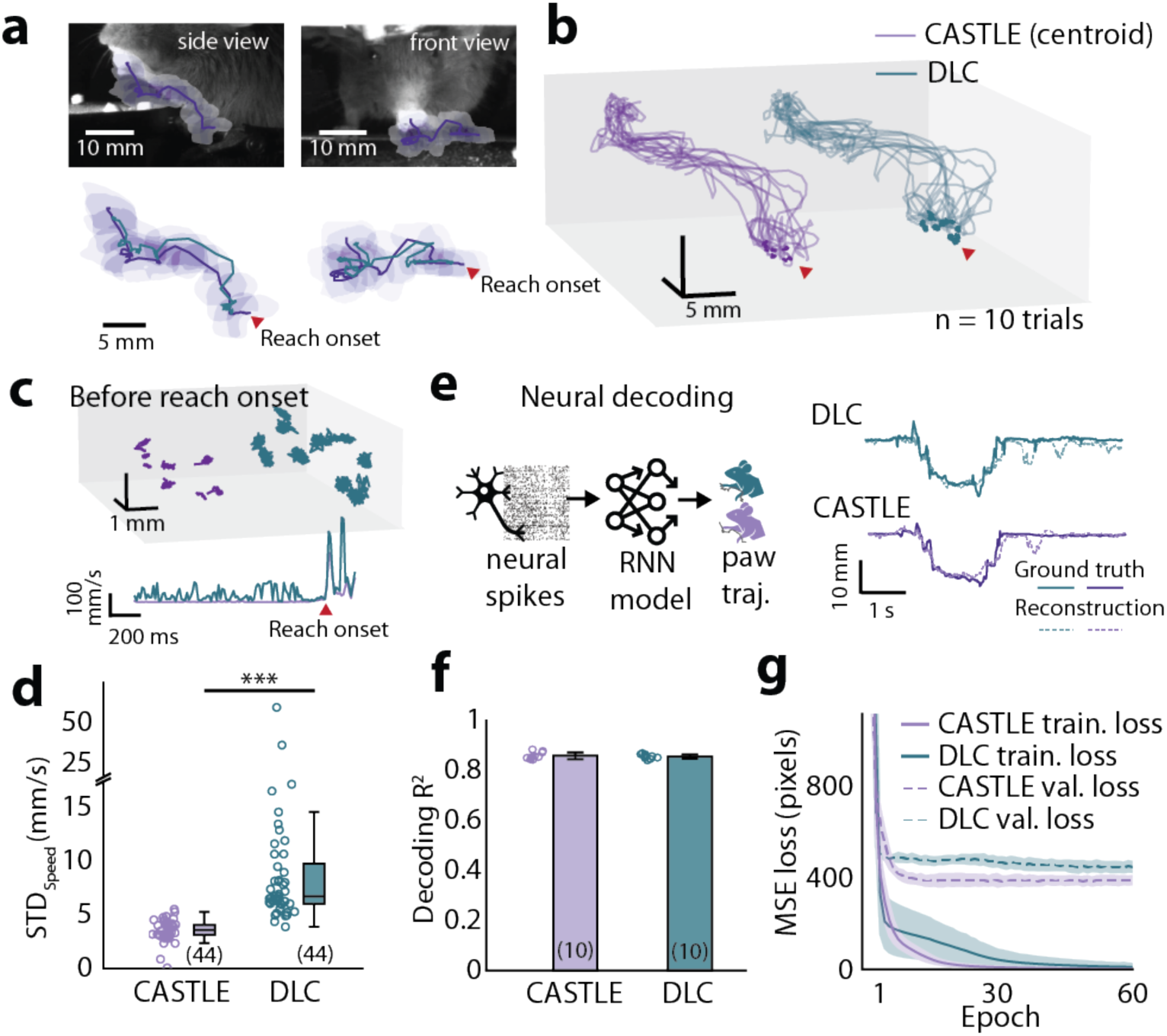
Comparative performance analysis between *CASTLE* and *DeepLabCut* (*DLC*) in tracking accuracy and neural decoding. **a.** Visualization of left paw trajectories from resting state to food grasping. Trajectories generated by *CASTLE* (purple lines), defined by connecting centroids of segmented ROI regions, are compared with *DLC*-derived trajectories (green-blue lines) using wrist-labeled tracking. **b.** Three-dimensional representation of ROI centroid trajectories across 10 independent trials. Darker segments denote trajectories before reach onset, highlighting greater displacement and higher noise levels in *DLC*-derived trajectories compared to *CASTLE*. **c-d.** Quantitative comparison of movement jitter measured as standard deviation (STD) of paw speed during resting states. *CASTLE* shows significantly lower jitter than *DLC* across 44 trials, as confirmed by statistical analysis (Wilcoxon signed-rank test, p < 0.001)**. e-g.** Neural decoding accuracy using simultaneously recorded neural spike data. A recurrent neural network (comprising two-layer GRUs and fully connected layers) trained with *CASTLE*-derived trajectories achieved comparable decoding performance for paw position (**f**) and lower mean squared error (MSE) loss during validation (**g**), indicating equivalent or superior performance relative to DLC-based keypoint tracking.

We further validated tracking performance by employing neural decoding as a benchmark. Accurate paw position tracking is crucial for interpreting the neural correlates of behavior, making neural decoding performance a suitable and biologically relevant measure for assessing tracking accuracy (see *Method details*). Neural decoding analyses using simultaneously recorded neural spiking activity from sensorimotor cortices indicated comparable decoding accuracy for paw positions (R^2^) between *CASTLE* and *DLC* (n = 10 independently trained models, *CASTLE*: R^2^ = 0.8554 ± 0.0135, *DLC*: R^2^ = 0.8524 ± 0.0086, student t-test, p = 0.5791 **Fig. 3e-f**). Importantly, models trained using *CASTLE*-derived trajectories demonstrated significantly lower mean squared error (MSE) during training and validation (**Fig. 3g**). In addition, similar tracking performance of *CASTLE* compared with *DLC* is also achievable in OFTs (**Extended Data Fig. 4**). Collectively, these findings underscore the practical utility, efficiency, and robustness of *CASTLE*, demonstrating its suitability for high-throughput, reliable behavioral studies; details on the required human engagement and computation time are provided in the *Methods* section.

### Behavioral classification in a mouse reach-and-grasp task using *CASTLE*

Given *CASTLE*’s advantages in efficiently capturing subtle behavioral dynamics without extensive manual labeling, we explored its capacity to identify detailed behavioral classes within complex reach-and-grasp tasks. These tasks present substantial analytical challenges due to the intricate and subtle variations in paw postures and movements^20^, which standard key-point labeling methods often fail to capture adequately. Specifically, typical key-point labeling methods, such as *DLC*, require detailed and extensive labeling of anatomical landmarks (e.g., finger joints or ankle positions) to accurately represent nuanced paw postures during grasping^9,21^. Such detailed labeling is not only highly time-consuming but also demands high-quality video data, significantly increasing costs and reducing practical usability. *CASTLE* leverages the visual foundation model *DINOv2* to directly encode comprehensive visual information from video frames, overcoming these limitations.

Initially, we manually defined five behavioral clusters: "on perch," "lifting," "grabbing," "at mouth," and "on stage" (**Fig. 4a, c**). *CASTLE* generated high-dimensional focused visual latent features from selected ROIs, encoding semantic visual information from front and side views (768 dimensions each, totaling 1536 dimensions; **Fig. 4a**). Movement kinematic features (position, speed, area) were extracted during segmentation and tracking but did not notably enhance classification quality (**Extended Data Fig. 5a**). Ablation studies confirmed the necessity of both “focused ROI extraction” and “visual latent features” for effective classification (**Extended Data Fig. 5b, c**).

**Figure 4.**
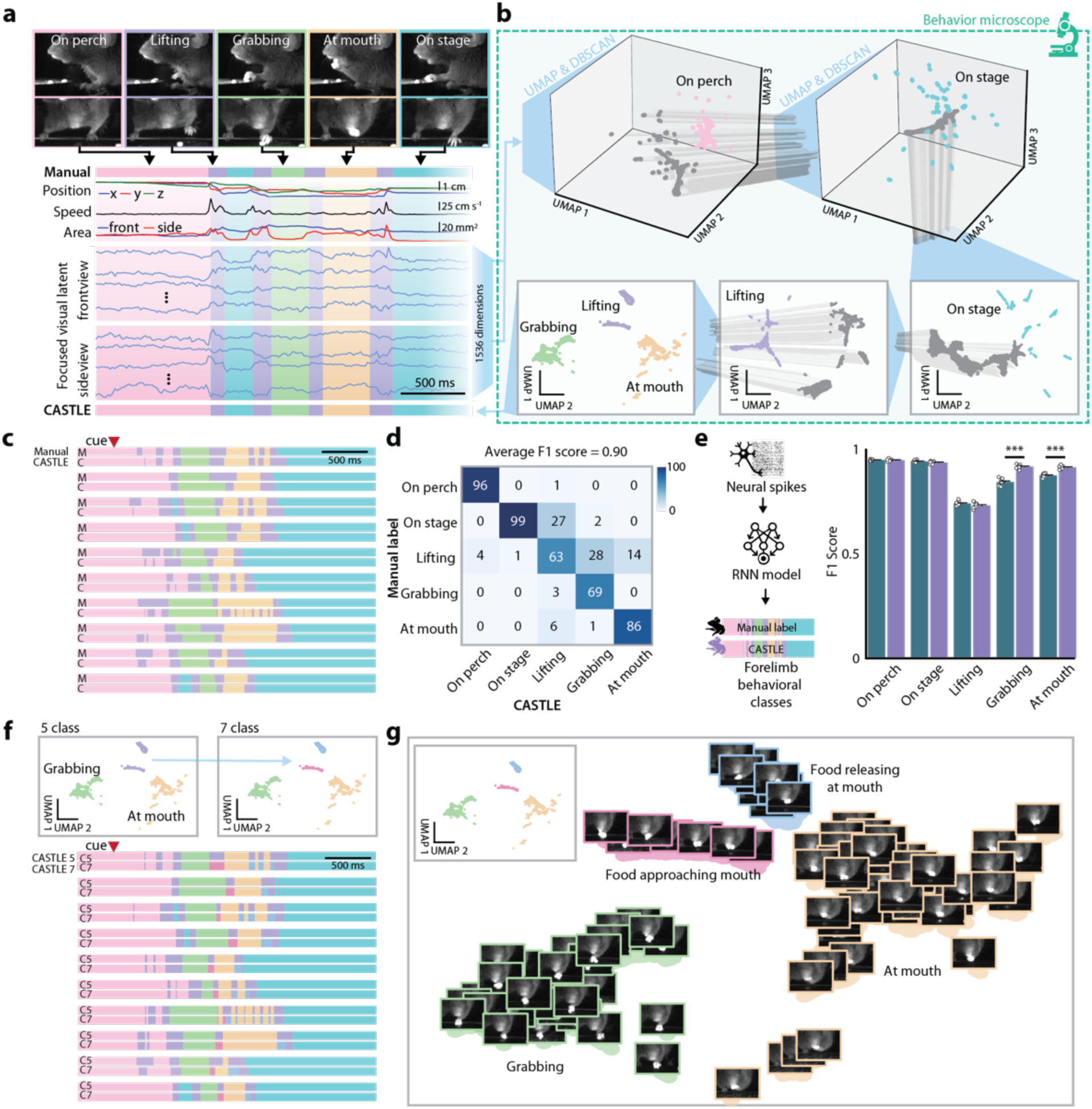
Extraction and hierarchical exploration of behavioral classes in a mouse reach-and-grasp task. **a.** Following the tracking of ROI trajectories, *CASTLE* employs *DINOv2* to extract focused visual latent representations from selected ROIs. These latent features represent semantic visual information within high-dimensional (768D per view) spaces, derived from both front and side views, totaling 1536 dimensions. Kinematic data including position, speed, and area are integrated alongside the visual latent features. Five clusters identified by *CASTLE* correspond directly to manually annotated behavioral phases during mouse reaching: "on perch," "lifting," "grabbing," "at mouth," and "on stage." **b.** Hierarchical clustering process in Latent Explorer, demonstrating iterative dimension reduction to simplify and visualize complex latent structures. In the initial stages, global structures were identified by employing parameters focusing on broad differences (see **Extended Data Fig. 2 and 3**). Subsequent stages progressively refined local structures by adjusting *UMAP* parameters, effectively isolating distinct behaviors. The hierarchical strategy enabled identification of subtle behavioral distinctions not initially anticipated by researchers (**Supplementary Video 4**). **c-d.** Reliability assessment of *CASTLE*-derived behavioral classifications compared to expert annotations, yielding an average F1 score of 0.90. Confusion matrix analysis indicated key discrepancies in classifications involving transitions among "Lift," "Grab," and "At mouth" behaviors. **e.** Neural activity recorded simultaneously was used to evaluate the classification performance of *CASTLE* versus expert annotations. A recurrent neural network (two-layer GRU with fully connected layers) was trained to predict behaviors from neural data across 36 trials. Over 10 independent training runs, there were no significant performance differences for the “on perch,” “on stage,” and “lifting” classes. However, performance on the “grabbing” and “at mouth” classes differed significantly, suggesting that the features defining these classes may be captured differently by the two labeling methods (n = 10, Wilcoxon rank-sum test p _on perch_ = 0.880, p _on stage_ = 0.226, p _lifting_ = 0.0963, p _grabbing_ = 0.000157, p _at mouth_ = 0.000157). **f-g.** *CASTLE* identified two distinct "Lift" behaviors overlooked by initial expert annotations: a transitional lift between "grabbing" and "at mouth," and a lift post "at mouth." Embedding visualizations illustrate how *CASTLE* clusters separate these subtle behaviors, extending researchers’ behavioral definitions beyond pre-established expectations, highlighting *CASTLE’s* utility in exploring and defining novel behavioral classes.

To systematically explore behavioral clusters, we applied a hierarchical clustering strategy in *CASTLE*’s “Behavior Microscope” (**Fig. 4b** and **Supplementary Video 3**). Iterative dimensionality reduction via *UMAP* first captured broad behavioral distinctions, subsequently refining local latent structures to isolate subtle differences (see **Extended Data Fig. 2 and 3a-d** for *UMAP* parameters consideration), similar to the delineation of skilled forelimb movements described previously^9,20,21^ (**Fig. 4b**).

We validated *CASTLE*-derived classifications against expert annotations, achieving a weighted average F1 score of 0.9015. Confusion matrix analysis showed discrepancies primarily involving transitions between "lifting," "grabbing," and "at mouth" behaviors, likely due to expert labeling variability and subtle behavioral transitions (**Fig. 4c-d**). To objectively assess these classifications, we employed neural decoding, as neural signals inherently reflect specific motor behaviors. Neural decoding models (a two-layer gated recurrent unit (GRU)^22^ with fully connected layers) trained on neural spiking activity across 36 trials demonstrated that *CASTLE*-derived behavior classifications significantly enhanced decoding accuracy for "grabbing" and "at mouth" compared to expert annotations ("grabbing", manual label: f1 score = 0.8459 ± 0.0143, *CASTLE*: R^2^ = 0.9193 ± 0.0085, Wilcoxon rank-sum test, p < 0.001; "At mouth ", manual label: R^2^ = 0.8770 ± 0.0082, *CASTLE*: R^2^ = 0.9162 ± 0.0080, Wilcoxon rank-sum test, p < 0.001), aligning closely with underlying neural representations (**Fig. 4e**). Notably, *CASTLE* identified two subtle behavioral variations overlooked by expert annotations: "food approaching mouth" (transitional lift between "grabbing" and "at mouth") and "food releasing at mouth" (occurring post "at mouth"). Visual embeddings distinctly separated these behaviors, demonstrating *CASTLE*’s potential in uncovering novel behavioral classes and providing deeper insights into skilled motor sequences (**Fig. 4f-g** and **Supplementary Video 4**).

### Comprehensive behavioral class identification in mouse open field tests (OFTs) using *CASTLE*

Considering the limitations of traditional OFT in automatically distinguishing detailed behavioral patterns, we leveraged the *CASTLE* framework to enhance behavioral classification. Traditional OFTs rely on basic metrics such as locomotion, exploratory activity, and spatial preferences, lacking the ability to automatically classify more complex behaviors like grooming and rearing^23,24^. Here we show that *CASTLE* significantly advances behavioral analysis by automating the precise identification of subtle behaviors such as grooming, supported rearing, and unsupported rearing without extensive manual annotation, thus improving sensitivity, accuracy, and efficiency.

We applied the *CASTLE* pipeline to analyze a 30-minute video recording of a mouse in an OFT environment at 30 frames per second (fps) (**Fig. 5a**), aiming for automated classification of detailed behaviors. To ensure precise behavioral representation, we removed irrelevant positional and directional features by cropping and aligning ROIs, a critical “focusing” step in *CASTLE* (**Fig. 5a**). ROIs for the body and tail-base were identified via Video Object Segmentation (VOS), from which angular metrics (angle, angular velocity) and focused visual latent features were extracted (**Fig.5b**). Directly using kinematic features, including speed, angular velocity, and area of ROI, for hierarchical latent exploration did not achieve clear behavioral classes (**Extended Data Fig. 5d**). Hierarchical latent exploration using sequential *UMAP* dimensionality reduction enabled effective classification of subtle behaviors from focused visual latent features (768 x 5 dimensions; **Fig.5c**). Initially, grooming behavior was distinctly separated using *UMAP* (time window = 5 frames, 167 ms). A subsequent two-stage *UMAP* approach identified nuanced differences between supported and unsupported rearing behaviors. Due to the high visual similarity between walking and immobility states, which could not be distinguished based on visual features alone, we applied a kinematic threshold of 4 mm/s (**Fig.5c** and **Supplementary Video 5**)^25^.

**Figure 5.**
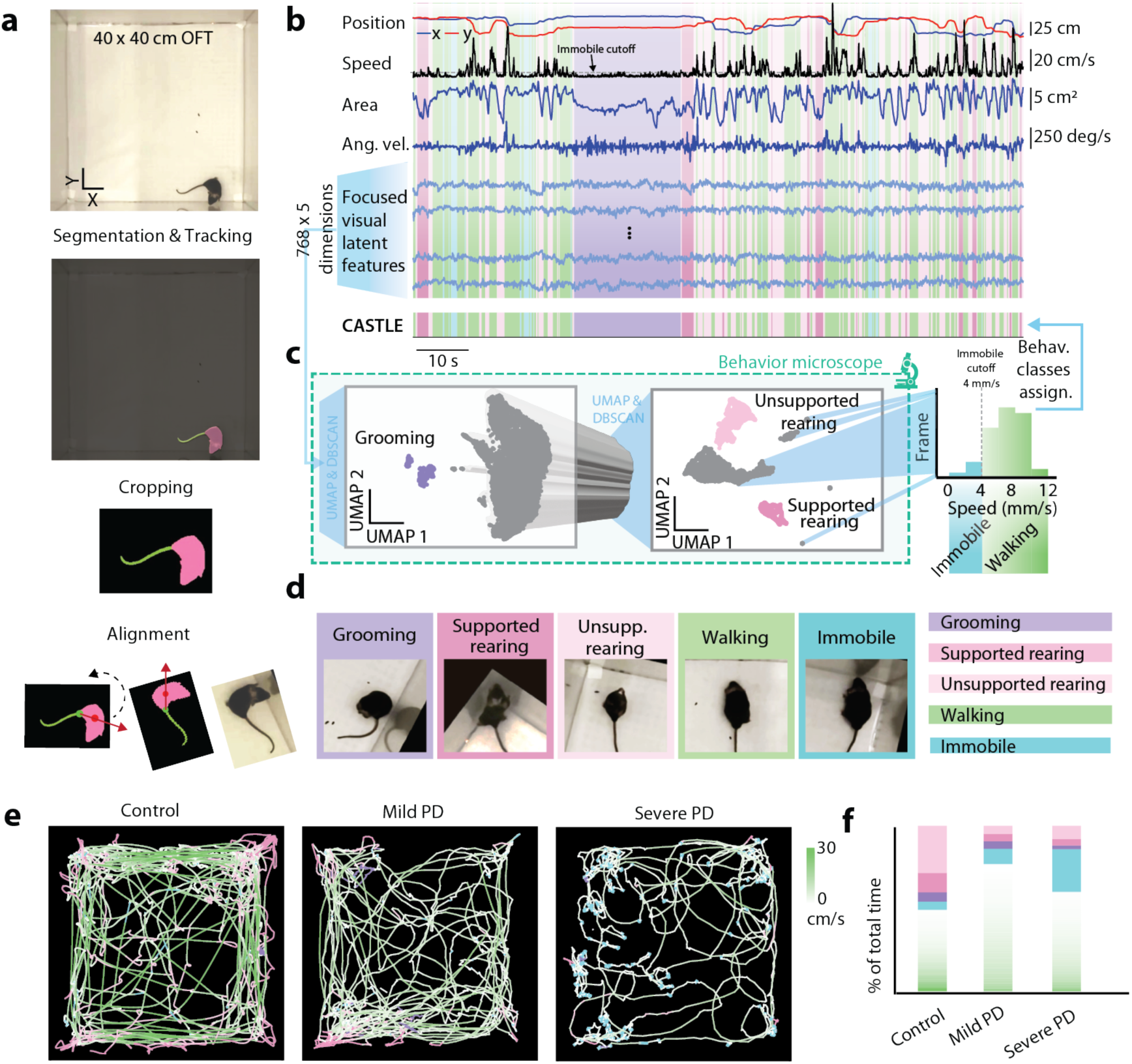
Behavioral classification using *CASTLE* in an OFT. **a.** Workflow illustrating the preprocessing steps for behavioral analysis using *CASTLE*. Initially, ROIs were identified using VOS. ROIs were cropped and aligned to remove spatial and directional bias, after which, focused visual latent features (*L_ROI_(t)*) were extracted using *DINOv2*, excluding irrelevant positional and directional information. **b.** Representative kinematic (position, speed, area, angular velocity, and 4 example visual latent features) and visual latent data traces derived from the mouse body and tail-base ROIs. Behavioral classes (grooming, supported rearing, unsupported rearing, walking, and immobile) were identified through latent exploration. **c.** Hierarchical latent exploration pipeline. An initial *UMAP* stage isolates grooming behavior. Subsequently, a 2-stage *UMAP* strategy delineates supported and unsupported rearing. Walking and immobile states are further distinguished using a kinematic speed threshold (4 mm/s). **d.** Representative video frames illustrating classified behaviors. **e-f.** Behavioral trajectories and analyses of control, mild Parkinson’s disease (PD), and severe PD mice. Walking speeds are visualized by color gradients, with darker colors indicating higher speeds. Severe PD mice exhibited significantly increased immobility.

*CASTLE* successfully classified five distinct behavioral classes: grooming, supported rearing, unsupported rearing, walking, and immobility (**Fig.5d**). Comparative analyses across control and mouse models of Parkinson’s disease (PD) — mild (acute MPTP-induced PD model) and severe (6-OHDA-induced hemi-PD model) — revealed distinct behavioral phenotypes (**Fig.5e**). Specifically, mild PD mice exhibited increased overall walking time but reduced occurrences of supported and unsupported rearing. Severe PD mice demonstrated increased immobility, reduced grooming, decreased rearing, and significantly slower walking speeds (**Fig.5f**). These findings highlight *CASTLE’s* robust ability to accurately characterize detailed behavioral phenotypes in both healthy and disease states, underscoring its value in high-throughput behavioral neuroscience research.

### Neutralizing location and orientation effects for improved classification

In developing *CASTLE*, we observed that appropriately focusing on ROIs is crucial to extract meaningful latent features for behavior classification. This observation implies that although VFMs capture abundant image information, much of it may be irrelevant or even detrimental for behavioral analysis. One particularly problematic factor is the animal’s orientation to the camera. Orientation changes can produce large shifts in the latent feature space that are unrelated to behavior *per se* (**Fig. 6a,b**). Orientation-sensitive features can sometimes be mitigated by physically aligning subjects in preprocessing (*e.g.*, always orienting the head in the same direction in each frame; **Fig. 5a**), but such alignment is often non-trivial for freely moving animals, especially for animals like *C. elegans*, or when video quality is limited. We therefore explored strategies to neutralize orientation in the latent features themselves.

**Figure 6.**
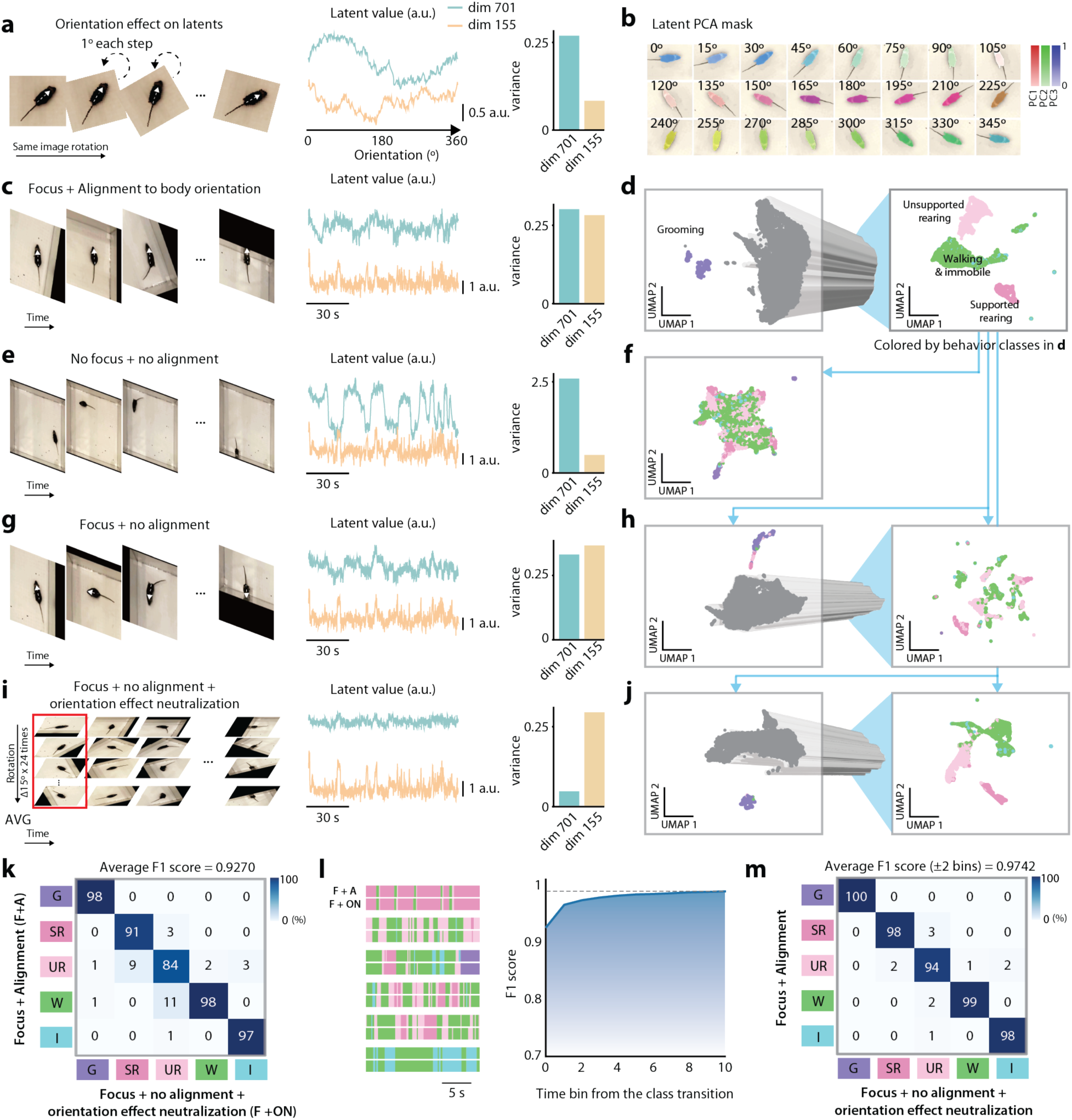
Effective extraction of focused latent features with orientation effect neutralization. **a.** Analysis of orientation sensitivity within focused visual latent features derived from *DINOv2*. The analysis demonstrates rotation-induced variations in latent values across dimensions by rotating a single mouse locomotion frame through 360 degrees. Results highlight dimensions with high orientation sensitivity (dimension 701) and those with minimal orientation dependence (dimension 155). **b.** Visualization of angular variations in focused visual latent features through RGB representation of the first three principal components (PC1-3) across different orientations, demonstrating the comprehensive influence of orientation effects across entire latent dimensions. **c-d.** Implementation of the baseline approach combining Focus and Alignment preprocessing to eliminate positional and orientation information. This method categorizes mouse behaviors into four distinct classes: grooming, supported rearing, unsupported rearing, and combined walking and immobile states. **e-f.** Latent exploration results from raw OFT video frames without preprocessing, demonstrating significant position-dependent clustering that obscures meaningful behavioral distinctions and limits effective behavioral classification. **g-h.** Latent exploration results following Focus preprocessing alone, which removes positional information but retains orientation dependencies. This approach shows improved clustering compared to raw data but still exhibits orientation-related artifacts that limit behavioral discrimination. **i-j.** Latent exploration results following Orientation Neutralization preprocessing, where averaged latent features (*L_ROI_(t)*) from multiple rotated frames effectively eliminate orientation dependencies while preserving behavioral distinctions for accurate classification. **k.** Performance comparison through weighted F1 scores between the baseline Focus and Alignment approach versus the Focus and Orientation Neutralization method, demonstrating equivalent classification accuracy without requiring reliable body orientation alignment. **l.** Temporal tolerance analysis examining F1 score performance across varying time bin tolerance levels. The improved accuracy with increased temporal tolerance validates that misclassifications are primarily transition-related differences. **m.** Confusion matrix visualization under ±2-time bin (±330 milliseconds) tolerance conditions, illustrating classification performance and error patterns between the alignment-based reference standard and Orientation Neutralization predictions.

As a first step, we quantified how rotations of the input frames affect the latent representation. We took example video frames and rotated them by various angles, then examined the *DINOv2* latent features (**Fig. 6a,b**). Qualitative inspection indicated that certain latent dimensions were highly sensitive to rotation, while others were relatively invariant (**Fig. 6a**). Comparing to classification with focused latent extraction and alignment (**Fig. 6c,d**), directly extract the latent features from original image frames revealed a strong clustering by absolute orientation and position rather than by behavior, which degraded the quality of behavioral clustering (**Fig. 6e,f**). Removing only positional information by introducing “focused” latent feature extraction partially reduced orientation-related variability, offering moderate improvements in behavioral clustering (**Fig. 6g-h**). We then implemented a more comprehensive rotation-averaging approach: for each video frame, we generated 24 rotated copies (every 15°) and averaged the latent embeddings across all these rotations. This approach dramatically reduced orientation-specific variance in the latent space and yielded much cleaner behavioral clusters (**Fig. 6i,j**). Using the orientation-neutralized latent features, *CASTLE* achieved high classification accuracy for all behaviors (average F1 scores > 0.9), with misclassifications largely limited to frames at the boundaries between behaviors (**Fig. 6j,k**). Analyses of transition probabilities and confusion (**Fig. 6l,m**) further confirmed that orientation-neutralization greatly improved the consistency and reliability of clustering—errors were minor and predominantly occurred at true behavior transition points. In summary, our results demonstrate that a rotation-based preprocessing strategy can robustly neutralize orientation effects in VFM-based latent features, thereby enhancing automated behavioral classification, especially for organisms or setups where consistent orientation alignment is infeasible.

### Behavioral classification of flies using *CASTLE*

Flies are widely used model organisms in behavioral neuroscience due to their well-defined behavioral repertoire, genetic tractability, and suitability for high-throughput screening^6,26,27^. Conventional behavioral studies in flies have relied heavily on manual annotation or supervised machine learning approaches, which require substantial training and user intervention^27^. Pose estimation methods like *DLC*^9^, *LEAP*^21^, and *SLEAP*^28^ have been developed to track fly movements precisely but require high-resolution video and extensive manual annotation for key-point labeling, making these methods resource-intensive and time-consuming^21^. To address these limitations, we extended the applicability of the *CASTLE* pipeline to automatically analyze and classify the complex behaviors of flies.

Using *CASTLE*, we analyzed behavior of flies in open-field test by sequentially performing background subtraction, segmentation, tracking, and orientation neutralization (**Fig. 7a**). Short 10-frame snippets of the flies’ focused latent features were embedded into a low-dimensional feature space via *UMAP* for unsupervised class discovery. A density-based clustering in this *UMAP* space yielded distinct behavioral clusters with minimal user input – no manual labeling or predefined behaviors were required. By eliminating a priori definitions of behaviors, *CASTLE*’s “Behavior Microscope” approach realizes balanced efficiency and interpretability in behavioral identification. Notably, whereas earlier unsupervised frameworks mapped hundreds of micro-behavioral “syllables” in flies^6^, our single-stage *UMAP* embedding strategy distilled the repertoire into a concise set of interpretable classes. In total, seven predominant behavioral classes emerged from the *CASTLE* analysis (**Fig. 7b**). Each of these classes corresponded to a distinct cluster in the *UMAP* behavior-space (**Fig. 7c**) and was verified by manual inspection of video segments.

**Figure 7.**
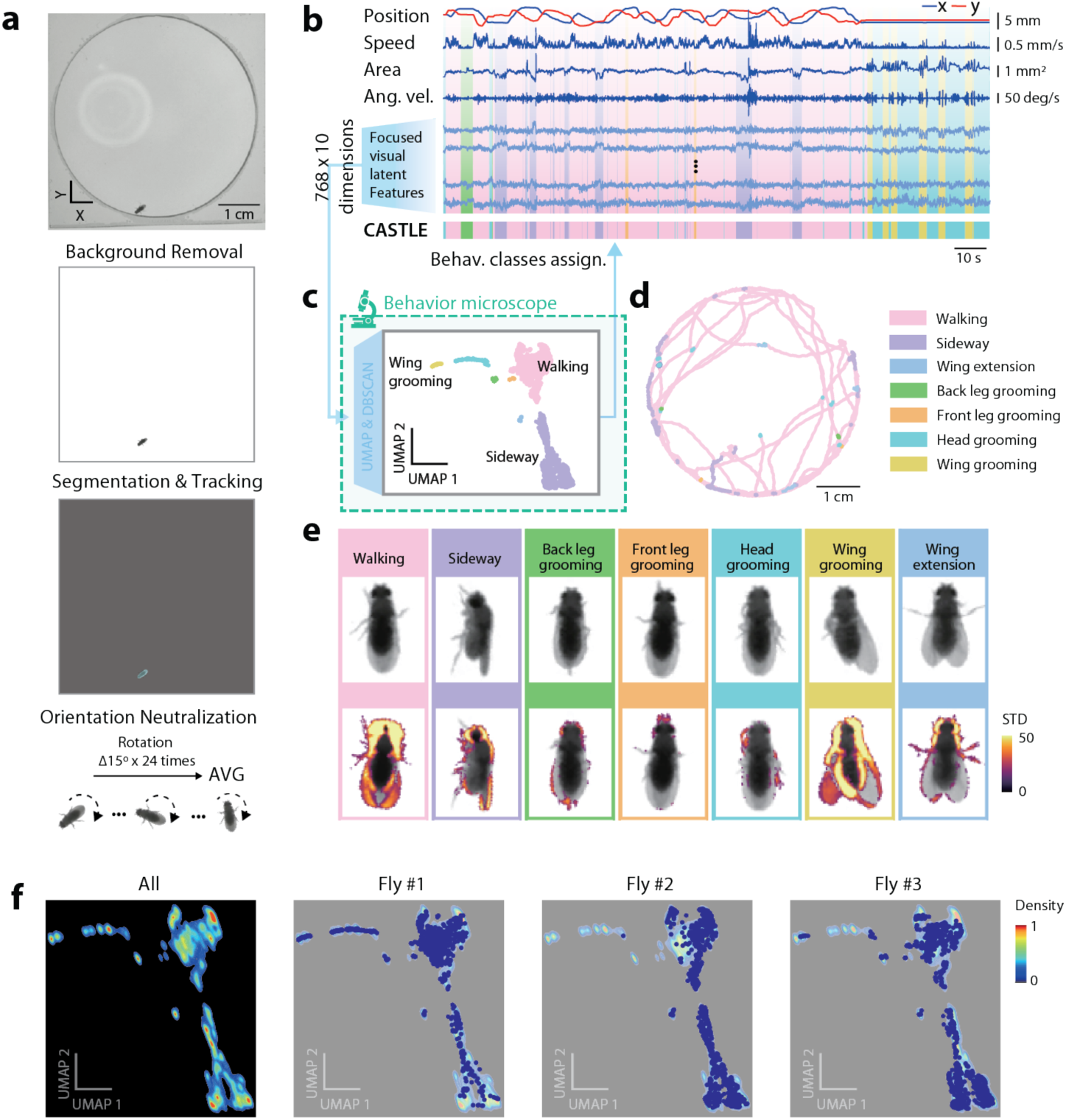
Application of *CASTLE* for behavioral classification in flies. **a.** Workflow for analyzing fly behavior from open-field test. The data include videos from 3 flies (3 minutes each at 60 fps). Steps include background removal, ROI segmentation and tracking, and orientation neutralization prior to feature extraction. **b-c.** Behavioral classes identified by *CASTLE*. A single-stage *UMAP* dimensionality reduction was applied to the focused visual latent features. Behaviors were clustered based on manual inspection and verification, using an optimal temporal binning (bin size = 10 frames, 6 Hz resolution) for analysis. **d.** Fly trajectories color-coded by their assigned behavioral clusters, demonstrating clear spatial segregation of different behaviors. **e.** Representative video frames and the standard deviation of pixel changes within a 15-frame interval highlighting diverse behaviors classified by *CASTLE*, such as grooming (front leg, back leg, head, and wing), walking, sideways movement, and wing extension. **f.** Density heatmaps showing the temporal density distribution of behaviors for three individual flies, revealing both common behavioral patterns and subject-specific differences in behavioral repertoires.

The clusters were further characterized by their quantitative feature profiles, including centroid position, translational speed, ROI area, angular velocity, and latent pose features. Characteristic kinematics were evident for several behaviors: for example, head grooming and wing grooming bouts showed pronounced changes in ROI area (due to limb or wing extension) and high angular velocity (calculated from the longest axis, which is significantly affected by wing movements). In contrast, front leg grooming lacked a unique signature in simple kinematic features and was distinguishable only in the high-dimensional visual latent space – reflecting the subtle, localized motion of the forelegs that *CASTLE*’s learned features could detect. Locomotor behaviors (walking and sideways walking) were associated with sustained movement and higher speed, whereas grooming and wing-extension behaviors occurred during brief pauses with little translational movement. We visualized the spatiotemporal distribution of these behaviors to confirm *CASTLE*’s classifications. The fly’s trajectory was color-coded by behavior classes (**Fig. 7d**), revealing that locomotion classes (walking modes) covered large areas of the arena, while grooming actions and wing extensions appeared as stationary or confined events (often occurring when the fly paused, frequently near the arena edges). Representative snapshots of the animal during each behavior are shown in **Fig. 7e**, highlighting distinctive postures — for instance, the extension of a single wing during wing grooming, or the forelegs touching the head during head grooming. Finally, we generated cluster-wise density maps of the flies’ behavioral classes (**Fig. 7f**). These maps indicate generalized and fly-specific differences in behavioral repertoires. In summary, *CASTLE* discovers meaningful structure in spontaneous fly behavior that aligns with known ethological motifs, without the extensive manual annotation required by supervised tracking tools like *SLEAP*^28^ or the complexity of earlier unsupervised mapping techniques^6^. This illustrates the power of combining video-based deep learning features with *UMAP-DBSCAN* clustering to catalog animal behavior in an objective, data-driven manner.

### Behavioral classification of *C. elegans* using *CASTLE*

Nematodes like *C. elegans* are staples of behavioral genetics, yet most locomotor analyses rely on curated ethograms, specialized single-worm (*WormTracker 2.0*^29^) or multi-worm trackers (*Multi-Worm Tracker* (*MWT*)^30^, *WormLab*, and *Tierpsy Tracker*^31^), or interactive classifiers that demand human supervision or require motorized stages and running in containerized environments. Against this backdrop, we apply *CASTLE* to freely-moving *C. elegans* and revealed that this pipeline can robustly distinguish both spontaneous locomotor patterns and optogenetically-evoked responses. After background subtraction, segmentation, tracking and orientation neutralization (24 rotations per mask; **Fig.8a**), five-frame snippets of the worm’s ROI were embedded via *UMAP* (**Fig.8b,c**). Density-based clustering revealed 5 discrete states: “Coiling” (tight, high-curvature curls), “Looping” (Ω-turns) (large body bends linked to reversals), “Local search” (area-restricted exploration with frequent reorientations), and extended search subdivided into “Forward (Global search)” and “Backward” movements (**Fig. 8c-e**). Direction was determined by comparing consecutive displacement vectors (**Fig. 8d**). Color-coded trajectory plots show that local search traces are clustered, global searches traverse the plate, reversals punctuate long runs, and coiling forms isolated dots (**Fig. 8f**). To test if optogenetic neural modulation affects locomotion behaviors, we stimulate an escape circuit with brief orange-light pulses (see *Methods*). Worms expressing a light-gated channel significantly shifted from sustained forward roaming into more looping and local search (N = 6, Looping: pre: 1.86 ± 0.81%, post: 10.82 ± 3.12%; Local search: pre: 13.4 ± 4.20%, post: 39.48 ± 5.39%; Global search, pre: 72.47 ± 7.77%, post: 31.40 ± 5.43%, all values expressed as mean ± SEM; Aligned Rank Transform (ART) ANOVA: stimulation × behavior interaction, p < 0.001; *post-hoc* Tukey-adjusted comparisons for Pre vs Post stimulation: looping p < 0.005, local search p < 0.001, global search p < 0.001, **Fig. 8g**). These shifts recapitulate the “pirouette” sequence documented in classical foraging studies and align with the forward–reverse–turn hierarchy mapped by neural circuit analyses^32–34^. Finally, skeleton-based body curvature patterns confirm that *CASTLE*’s classification outputs correspond to biomechanically distinct locomotor modes (**Fig. 8h** and **Extended Data Fig. 6**).

**Figure 8.**
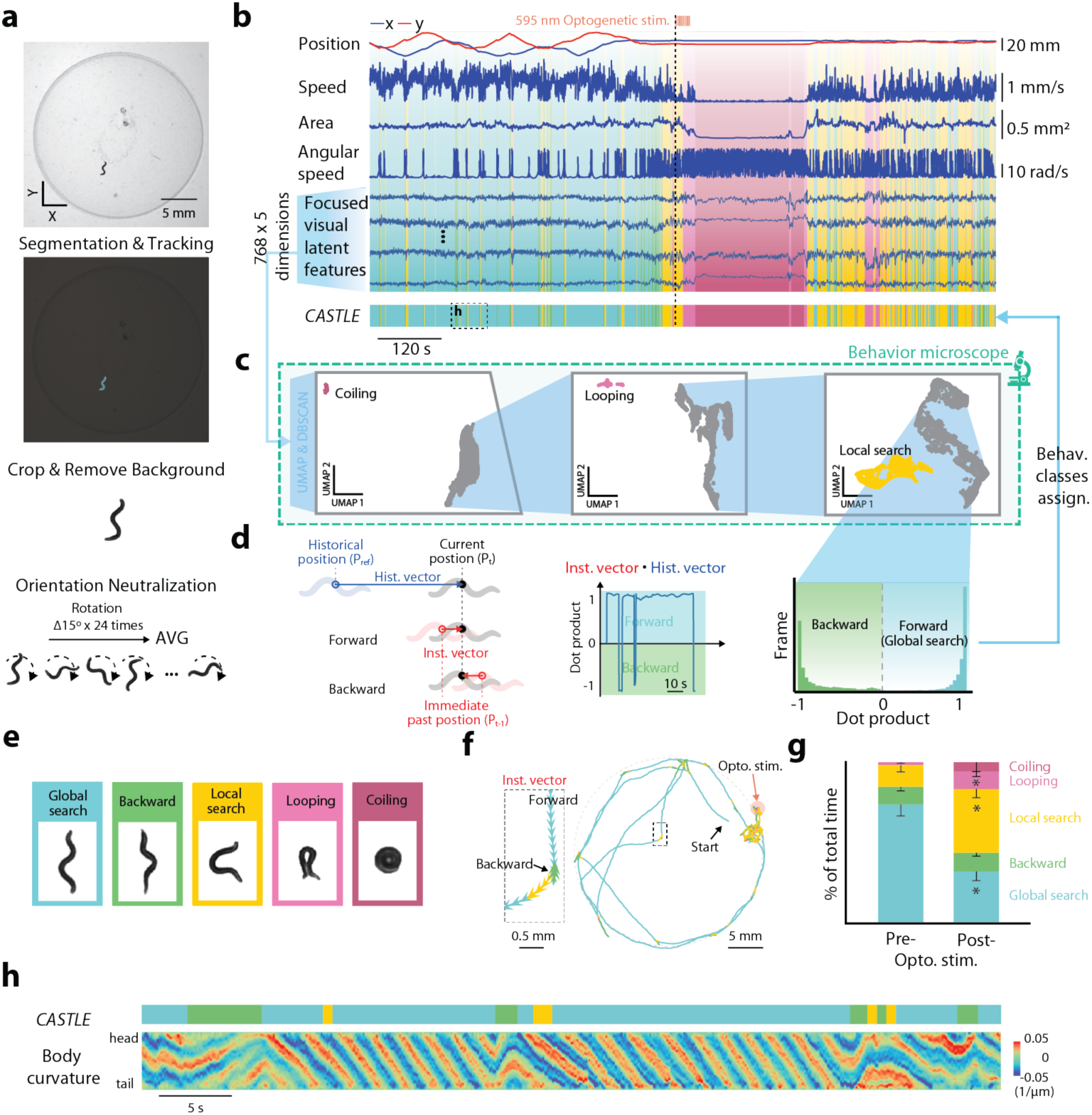
Application of *CASTLE* for behavioral classification in *C. elegans*. **a.** Workflow for analyzing six 20-minute *C. elegans* open-field videos (7.5 fps). The pipeline includes ROI segmentation and tracking, background removal to eliminate locomotion trails, and orientation neutralization to remove directional bias prior to focused visual latent feature extraction. Integrated visualization displaying traditional kinematic information (xy coordinates, speed, ROI areas, angular speed), focused visual latent features (*L_ROI_(t)*), and behavioral classes identified by *CASTLE*. The correlation between kinematic parameters and dynamics of focused visual latent representations are illustrated how focused visual latent features effectively capture behavioral distinctions that translate into discrete behavioral classifications. **c.** Latent exploration employing a three-stage hierarchical clustering approach. Step 1: Initial *UMAP* dimensionality reduction (*n_neighbors* = 30, *n_components* = 2, bin size = 5) successfully isolated the "coiling" behavioral cluster (red). Step 2: 2-stage *UMAP* architecture (*n_neighbors* = (300, 100), *n_components* = (5, 2)) identified and separated the "looping" behavioral cluster (pink). Step 3: Applying identical parameters to the remaining data distinguished the "local search" cluster. **d**. Since global search forward and backward movements exhibit similar morphological characteristics preventing visual distinction, directional classification utilizes positional trajectory analysis incorporating historical location, immediate past position, and current position to determine movement directionality. Algorithm for computing directional dot products to distinguish forward from backward locomotion, with displacement threshold D set at 2.5 mm. **e.** Representative video frames illustrating diverse behaviors classified by *CASTLE*, including global search, local search, backward movement, coiling and looping behaviors. **f.** *C. elegans* trajectories color-coded by their assigned behavioral cluster, demonstrating clear spatial segregation and characteristic movement patterns for different behavioral categories. The inset shows a trajectory segment that includes backward movement. **g**. Quantitative comparison of behavioral state proportions before and after optogenetic stimulation (N = 6, error bars indicate SEM). Statistical analysis was performed using Aligned Rank Transform (ART) ANOVA for repeated measures, revealing a significant stimulation × behavior interaction (p < 0.001). Post-hoc comparisons with Tukey adjustment demonstrated significant pre-post stimulation differences for looping (p < 0.01), local search (p < 0.001), and global search (p < 0.001) behaviors. **h.** Validation of *CASTLE*-identified behavioral classes through comparison with *C. elegans* body curvature dynamics. Curvature analysis confirms that *CASTLE* behavioral classifications correspond to distinct biomechanical signatures, providing independent verification of the *CASTLE* behavioral categorization through established postural analysis methods.

Notably, all six worms expressed these behaviors with pronounced individual differences (**Extended Data Fig. 7a**). Most worms spent most of their time roaming (global search forward) or performing local searches in control condition, while looping and coiling were observable after optogenetic stimulation. Some worms never coiled in the 20-min recording (**Extended Data Fig. 7b-e**); one produced multiple coils, especially immediate after optogenetically evoked escape responses. Despite this variability, the underlying repertoire was consistent across individuals. *CASTLE* provides an alternative to specialized single worm trackers. It yields posture-aware classifications with the interpretability of curated ethograms and delivers a unified framework for high-throughput phenotyping across species.

## Discussion

*CASTLE* combines zero-shot segmentation (*SAM*^15^), robust mask tracking (*DeAOT*^16^) and foundation-model vision (*DINOv2*^17^) to create posture-specific, orientation-neutral latent spaces that are hierarchically clustered into a multiscale ethogram via our interactive “Behavior Microscope” interface. By removing the need for labor-intensive annotation, task-specific retraining, and common bottlenecks for current methods, *CASTLE* enables rapid, training-free behavioral analysis while ensuring the results are interpretable and biologically validated.

### Relation to existing behavioral-classification tools

We position *CASTLE* within three common modules of behavioral pipelines to clarify feature differences and use-cases: (i) segmentation/tracking, (ii) image-information encoding, and (iii) clustering and classification. For segmentation and tracking, keypoint estimators such as *DLC*^9^, *Lightning Pose*^10^, and *SLEAP*^28^ output sparse landmarks that are ideal for kinematic readouts and 3D reconstructions when anatomy is visible, but they require labeled training data and can jitter or fail under occlusion. Depth/silhouette approaches, exemplified by *MoSeq*’s fixed top-down arenas, yield stable whole-body outlines yet are sensor- and geometry-dependent and often miss distal detail^35^. *CASTLE* uses dense mask tracking via VOS to preserve the full pixel footprint of each ROI without task-specific training, and keeps identity handling external and modular.

For image-information encoding, pose-derived features are compact and interpretable, but they are bounded by the chosen skeleton. Typical implementations derive kinematics or pose dynamics from tracked landmarks, for example *DLC*^9^, *Lightning Pose*^10^, and *SLEAP*^28^, *DeepPoseKit*^36^, *DeepFly3D*^37^, and *Keypoint*-*MoSeq*^38^. Silhouette or pixel embeddings capture global shape without landmarks, yet they can entangle orientation and view. Examples include aligned-pixel and depth representations used to map *Drosophila* behavior^6,39,40^ and depth-frame embeddings in *MoSeq*^35^. Learned pixel encoders can be highly discriminative, although supervised and self-supervised variants usually require retraining or curated clips when domains shift, as in *Selfee*^41^, *LabGym*^42^*,and DeepEthogram*^43^, and *DeepAction*^44^. *CASTLE* instead reuses a general-purpose visual foundation model^11,17^ and focuses it on ROIs, with optional rotation averaging to reduce orientation sensitivity. The trade-offs are reduced immediate interpretability relative to kinematics and a dependence on the priors of the chosen foundation model.

For clustering and classification, unsupervised pipelines such as *B-SOiD* characterize behavior as clusters of short-duration states at a chosen resolution^7^. Generative sequence models such as *MoSeq*^35^ and *Keypoint*-*MoSeq*^38^ jointly segment states and describe how they tend to progress over time, including typical transitions and dwell times. Supervised recognizers, *e.g.*, *JAABA*^27^, *SimBA*^45^, *VAME*^46^, and *DeepEthogram*^43^, deliver predefined labels once trained. *CASTLE* implements an interactive hierarchical alternative, the “Behavior Microscope”, which moves from broad structure to fine motifs with direct video verification. If needed, downstream models can be applied to *CASTLE* labels or features to quantify transition patterns and durations.

### Complementarity rather than exclusivity

These modules can be mixed and matched: keypoints from *DLC*^9^ and *SLEAP*^28^ can be clustered by *B-SOiD*^7^ or modeled by *Keypoint*-*MoSeq*^38^ to study how states unfold over time. Conversely, *CASTLE*’s ROI-focused latent features can be used with other clustering algorithms or with sequence models to analyze transition patterns and state durations. For multi-animal scenes, identity-aware trackers can provide per-animal masks that are then encoded and explored with *CASTLE*’s latent workflow. In practice, the optimal pipeline is determined by available labels and sensors, the required outputs such as coordinates, state labels, or transition statistics, and the degree of domain shift expected across experiments. This three-module framing emphasizes that *CASTLE* is one particular instantiation with dense ROI masks, training-free ROI-focused encoders, and hierarchical interactive clustering, making different trade-offs than keypoint-first or supervised action-recognition pipelines. It also highlights clear avenues for integration (e.g., coupling *CASTLE* latent features with sequence models) and for future benchmarking in multi-animal and 3D contexts.

### Limitations of *CASTLE*

Multi-animal tracking and analysis methods^28,47,48^, such as multi-animal *DLC*^48^, *SLEAP*^28^, *AlphaTracker*^47^, extend supervised pose estimation to social behavior settings. In the current work, we validate *CASTLE* only on single-animal datasets (flies and worms analyzed one per video). Because *CASTLE* is built on category-agnostic segmentation and per-ROI embeddings rather than a fixed skeleton, the architecture is intrinsically compatible with multi-animal scenes once distinct masks are available; benchmarking this extension against *DLC*^48^, *SLEAP*^28^ and *AlphaTracker*^47^ is a clear next step.

*CASTLE* also shares limitations typical of unsupervised, vision-based systems. First, despite zero-shot segmentation, users still seed masks and occasionally correct drift: usually a handful of frames per hour of video. Second, orientation neutralization improves clustering but increases inference time. This feature is improvable by reducing averaged angles per frame. Third, the latent space inherits biases of image-patch-based *DINOv2*^17^; under-represented body parts (*e.g.*, whisker or individual fly leg) may be harder to resolve. Finally, temporal context is incorporated by concatenating frame embeddings; behaviors that hinge on long-range sequence order may benefit from video-native foundation models.

### Future directions for *CASTLE*

Several extensions are natural. (i) Exchange models with state-of-the-art VFMs, such as *SAM 2*^49^ and *DINOv3*^50^. (ii) Video-native VFMs, *e.g.* V-JEPA 2^51^, could encode motion directly and support joint embeddings with audio or neural signals, tightening the link between latent clusters and underlying computations. (iii) Language grounding— attaching concise natural-language descriptors^14^ to clusters—could close the loop between automated discovery and human-readable ethograms. (iv) Closed-loop control: integrating *CASTLE*’s low-latency labels with experimental hardware would enable state-contingent stimulation within tens of milliseconds. (v) Social behavior: extending segmentation/tracking to multiple interacting animals will allow benchmarking against *SLEAP*^28^ and *AlphaTracker*^47^ in identity-aware settings.

## Conclusion

In summary, *CASTLE* occupies a distinctive middle-ground niche: as training-free and discovery-oriented as *B-SOiD*^7^ or *MoSeq*^35^, yet as spatially precise as *DLC*^9^, as video-aware as VFMs, and potentially scalable to social settings once multi-mask tracking is enabled. By exchanging tailored training for foundation-model vision and layering a multiscale clustering engine on top, *CASTLE* delivers a turnkey, versatile, cross-species solution that recovers expert-level annotations, uncovers hidden motifs and scales from limb dynamics to whole-body behavior. As foundation models advance, *CASTLE* should remain an adaptable platform for high-throughput, objective behavioral phenotyping in neuroscience and computational ethology.

## Acknowledgments

The authors thank Dr. Kuo-Hua Huang, Dr. Yi-Ping Hsueh, Dr. Yu Tsao, Dr. Jun Ding, and members of Wu laboratory for helpful discussions. This study was funded by grants from the Academia Sinica I-AI-A Grant AS-IAIA-114-L01 (Y.-W.W. and C.-L.H.); the Ministry of Science and Technology and National Science and Technology Council, Taiwan, NSTC 114-2321-B-001-005 (Y.-W.W.), NSTC 113-2321-B-001-012 (Y.-W.W.), NSTC 112-2321-B-001-007 (Y.-W.W.), MOST 111-2321-B-001-011 (Y.-W.W.), MOST 110-2321-B-001-012 (Y.-W.W.); Institute of Molecular Biology, Academia Sinica SPP (2024) and SPP (2025).

## Author contributions

Y.-S.L. and Y.-W.W. designed the analysis pipelines and the experiments. Y.-S.L. established the analysis pipeline and the GUI. Y.-S.L. performed the data analysis for behavioral classifications and neural decoding. Y.-S.L., H.-Y.Y, Y.-T.H, and B.-S.W.,. performed OFT of mice. Y.-S.L., Y.-W.W. and Y.-F.C performed reach-and-grasp test of mice. J.S. and S.L. performed OFT of flies. J.-B.Y. and C.-H.C. performed OFT of *C. elegans*. Y.-W.W. and Y.-S.L. made the figures. Y.-S.L., C.-L.H., and Y.-W.W. wrote the manuscript with contribution of all coauthors. Y.-W.W. conceived the study and supervised the research.

## References

1 Krakauer, J. W., Ghazanfar, A. A., Gomez-Marin, A., MacIver, M. A. & Poeppel, D. Neuroscience needs behavior: correcting a reductionist bias. Neuron 93, 480–490 (2017).

2 Guo, J.-Z. et al. Cortex commands the performance of skilled movement. Elife 4, e10774 (2015).

3 Brown, A. E. & De Bivort, B. Ethology as a physical science. Nature Physics 14, 653–657 (2018).

4 Han, Y. et al. Multi-animal 3D social pose estimation, identification and behaviour embedding with a few-shot learning framework. Nature Machine Intelligence 6, 48–61 (2024).

5 Marks, M. et al. Deep-learning-based identification, tracking, pose estimation and behaviour classification of interacting primates and mice in complex environments. Nature machine intelligence 4, 331–340 (2022).

6 Berman, G. J., Choi, D. M., Bialek, W. & Shaevitz, J. W. Mapping the stereotyped behaviour of freely moving fruit flies. Journal of The Royal Society Interface 11, 20140672 (2014).

7 Hsu, A. I. & Yttri, E. A. B-SOiD, an open-source unsupervised algorithm for identification and fast prediction of behaviors. Nature communications 12, 5188 (2021).

8 Goodwin, N. L. et al. Simple Behavioral Analysis (SimBA) as a platform for explainable machine learning in behavioral neuroscience. Nature Neuroscience, 1–14 (2024).

9 Mathis, A. et al. DeepLabCut: markerless pose estimation of user-defined body parts with deep learning. Nature neuroscience 21, 1281–1289 (2018).

10 Biderman, D. et al. Lightning Pose: improved animal pose estimation via semi-supervised learning, Bayesian ensembling and cloud-native open-source tools. Nature Methods, 1–13 (2024).

11 Awais, M. et al. Foundational models defining a new era in vision: A survey and outlook. arXiv 2023. arXiv preprint arXiv:2307.13721 (2023).

12 Zheng, C. et al. in 2024 IEEE International Conference on Big Data (BigData). 1909–1918 (IEEE).

13 Sun, J. J. et al. Video Foundation Models for Animal Behavior Analysis. bioRxiv, 2024.2007.2030.605655 (2024).

14 Xu, T., et al. MouseGPT: A Large-scale Vision-Language Model for Mouse Behavior Analysis. arXiv preprint arXiv:2503.10212 (2025).

15 Kirillov, A. et al. in Proceedings of the IEEE/CVF International Conference on Computer Vision. 4015–4026.

16 Yang, Z. & Yang, Y. Decoupling features in hierarchical propagation for video object segmentation. Advances in Neural Information Processing Systems 35, 36324–36336 (2022).

17 Oquab, M., et al. Dinov2: Learning robust visual features without supervision. arXiv preprint arXiv:2304.07193 (2023).

18 McInnes, L., Healy, J. & Melville, J. Umap: Uniform manifold approximation and projection for dimension reduction. arXiv preprint arXiv:1802.03426 (2018).

19 Ester, M., Kriegel, H.-P., Sander, J. & Xu, X. in kdd. 226–231.

20 Guo, J. Z. et al. Cortex commands the performance of skilled movement. Elife 4, e10774 (2015). 10.7554/eLife.10774

21 Pereira, T. D. et al. Fast animal pose estimation using deep neural networks. Nature methods 16, 117–125 (2019).

22 Chung, J., Gulcehre, C., Cho, K. & Bengio, Y. Empirical Evaluation of Gated Recurrent Neural Networks on Sequence Modeling. arXiv:1412.3555 (2014). <https://ui.adsabs.harvard.edu/abs/2014arXiv1412.3555C>.

23 Seibenhener, M. L. & Wooten, M. C. Use of the Open Field Maze to measure locomotor and anxiety-like behavior in mice. J Vis Exp, e52434 (2015). 10.3791/52434

24 Walsh, R. N. & Cummins, R. A. The Open-Field Test: a critical review. Psychol Bull 83, 482–504 (1976).

25 Sanders, T. H. & Jaeger, D. Optogenetic stimulation of cortico-subthalamic projections is sufficient to ameliorate bradykinesia in 6-ohda lesioned mice. Neurobiology of disease 95, 225–237 (2016).

26 Dankert, H., Wang, L., Hoopfer, E. D., Anderson, D. J. & Perona, P. Automated monitoring and analysis of social behavior in Drosophila. Nature methods 6, 297–303 (2009).

27 Kabra, M., Robie, A. A., Rivera-Alba, M., Branson, S. & Branson, K. JAABA: interactive machine learning for automatic annotation of animal behavior. Nature methods 10, 64–67 (2013).

28 Pereira, T. D. et al. SLEAP: A deep learning system for multi-animal pose tracking. Nature methods 19, 486–495 (2022).

29 Vedantham, K. et al. Track-A-Worm 2.0: A Software Suite for Quantifying Properties of C. elegans Locomotion, Bending, Sleep, and Action Potentials. bioRxiv (2024). 10.1101/2024.09.12.612524

30 Swierczek, N. A., Giles, A. C., Rankin, C. H. & Kerr, R. A. High-throughput behavioral analysis in C. elegans. Nat Methods 8, 592–598 (2011). 10.1038/nmeth.1625

31 Javer, A. et al. An open-source platform for analyzing and sharing worm-behavior data. Nat Methods 15, 645–646 (2018). 10.1038/s41592-018-0112-1

32 Kato, S. et al. Global brain dynamics embed the motor command sequence of Caenorhabditis elegans. Cell 163, 656–669 (2015). 10.1016/j.cell.2015.09.034

33 Flavell, S. W., Raizen, D. M. & You, Y. J. Behavioral States. Genetics 216, 315–332 (2020). 10.1534/genetics.120.303539

34 Huo, J. et al. Hierarchical behavior control by a single class of interneurons. Proc Natl Acad Sci U S A 121, e2410789121 (2024). 10.1073/pnas.2410789121

35 Markowitz, J. E. et al. The striatum organizes 3D behavior via moment-to-moment action selection. Cell 174, 44–58. e17 (2018).

36 Graving, J. M. et al. DeepPoseKit, a software toolkit for fast and robust animal pose estimation using deep learning. Elife 8 (2019). 10.7554/eLife.47994

37 Günel, S. et al. DeepFly3D, a deep learning-based approach for 3D limb and appendage tracking in tethered, adult Drosophila. Elife 8, e48571 (2019).

38 Weinreb, C. et al. Keypoint-MoSeq: parsing behavior by linking point tracking to pose dynamics. Nature Methods 21, 1329–1339 (2024).

39 Berman, G. J., Bialek, W. & Shaevitz, J. W. Predictability and hierarchy in Drosophila behavior. Proceedings of the National Academy of Sciences 113, 11943–11948 (2016).

40 Todd, J. G., Kain, J. S. & de Bivort, B. L. Systematic exploration of unsupervised methods for mapping behavior. Physical biology 14, 015002 (2017).

41 Jia, Y. et al. Selfee, self-supervised features extraction of animal behaviors. Elife 11, e76218 (2022).

42 Hu, Y., et al. LabGym: Quantification of user-defined animal behaviors using learning-based holistic assessment. Cell Reports Methods 3 (2023).

43 Bohnslav, J. P. et al. DeepEthogram, a machine learning pipeline for supervised behavior classification from raw pixels. Elife 10 (2021). 10.7554/eLife.63377

44 Harris, C., Finn, K. R., Kieseler, M.-L., Maechler, M. R. & Tse, P. U. DeepAction: a MATLAB toolbox for automated classification of animal behavior in video. Scientific Reports 13, 2688 (2023).

45 Goodwin, N. L. et al. Simple Behavioral Analysis (SimBA) as a platform for explainable machine learning in behavioral neuroscience. Nat Neurosci 27, 1411–1424 (2024). 10.1038/s41593-024-01649-9

46 Luxem, K. et al. Identifying behavioral structure from deep variational embeddings of animal motion. Communications Biology 5, 1267 (2022).

47 Chen, Z. et al. AlphaTracker: a multi-animal tracking and behavioral analysis tool. Frontiers in Behavioral Neuroscience 17, 1111908 (2023).

48 Lauer, J. et al. Multi-animal pose estimation, identification and tracking with DeepLabCut. Nature Methods 19, 496–504 (2022).

49 Ravi, N., et al. Sam 2: Segment anything in images and videos. arXiv preprint arXiv:2408.00714 (2024).

50 Siméoni, O., et al. Dinov3. arXiv preprint arXiv:2508.10104 (2025).

51 Assran, M. et al. V-jepa 2: Self-supervised video models enable understanding, prediction and planning. arXiv preprint arXiv:2506.09985 (2025).

