## Supplementary Information for "CASTLE: a training-free foundation-model pipeline for cross-species behavioral classification"

### Extended Data figures and legends

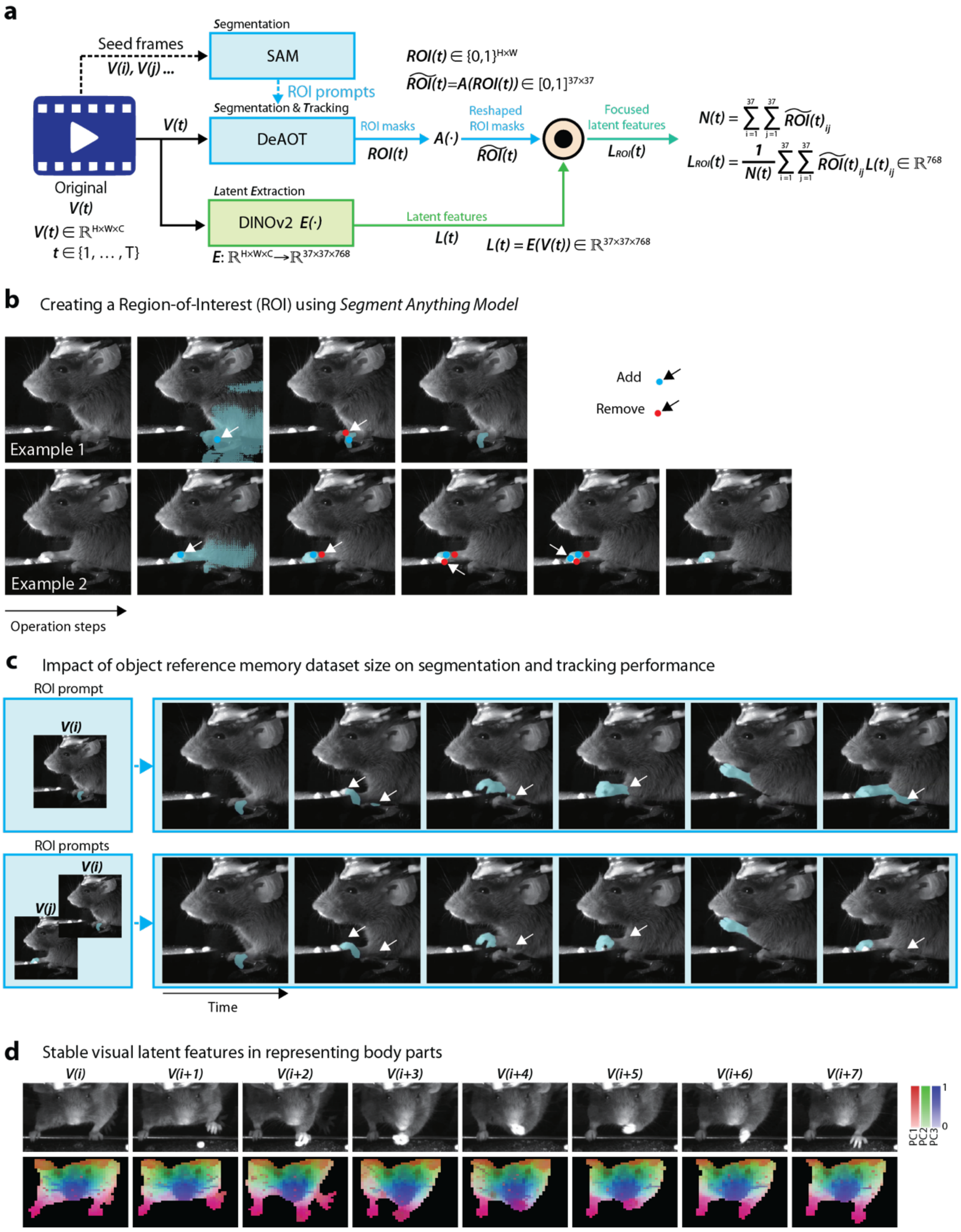

**Extended Data Figure 1. Interactive ROI segmentation, video object tracking, and broadly-focused patch latent feature visualization in the *CASTLE* pipeline**

**a.** Workflow for computing focused visual latent features from video frames. The pipeline transforms raw video frames (H×W×C dimensions) through ROI segmentation and *DINOv2* feature extraction. ROI masks ($ROI(t)$) obtained from *DeAOT* are reshaped to match *DINOv2*'s 37×37 patch grid, generating weight matrices $\widetilde{ROI}(t)$ where each element represents the proportion of patch-ROI overlap. The focused visual latent $L_{ROI}(t)$ is computed via weighted averaging, combining patch latents $L(t)_{ij}$ with corresponding ROI weights and normalizing by $N(t)$ to account for variable ROI sizes. This process yields a 768-dimensional focused visual latent feature per ROI per frame, creating a time-series dataset optimized for behavioral analysis. **b.** Interactive ROI creation using an interactive segmentation model. The user refines the mask by clicking on the image (white arrows indicate click locations). Blue points expand the selection (add), while red points contract it (remove). The sequence from left to right shows how the model's predicted mask updates with each click. Typically, within a few clicks, the model accurately identifies the user's intended body part. Representative examples illustrate the steps for annotation of a single frame. **c.** Impact of reference memory on ROI tracking using a video object segmentation, such as *DeAOT* models. The models use these initial, interactively defined ROIs as input prompts to predict ROIs in subsequent, unannotated frames. If only a single ROI is provided as the initial prompt (top scenario), tracking errors can occur during challenging movements. For instance, when a mouse lifts one hand, the mask might incorrectly track the other, leading to cascading errors. However, providing multiple ROIs as initial prompts (bottom scenario) enhances the models' contextual understanding, enabling it to maintain accurate tracking on the correct body part even when the animal makes complex movements like lifting a hand. White arrows indicate the errors that were corrected by this approach **d.** Visualization of mouse patch latent features using PCA (**Supplementary Video 1**). The top three principal components (PCs), derived from a global analysis of all mouse-related patch latent features, are mapped to RGB color channels. This reveals that distinct body parts are the primary features encoded, with their color representation remaining consistent for the same body part across different video frames.

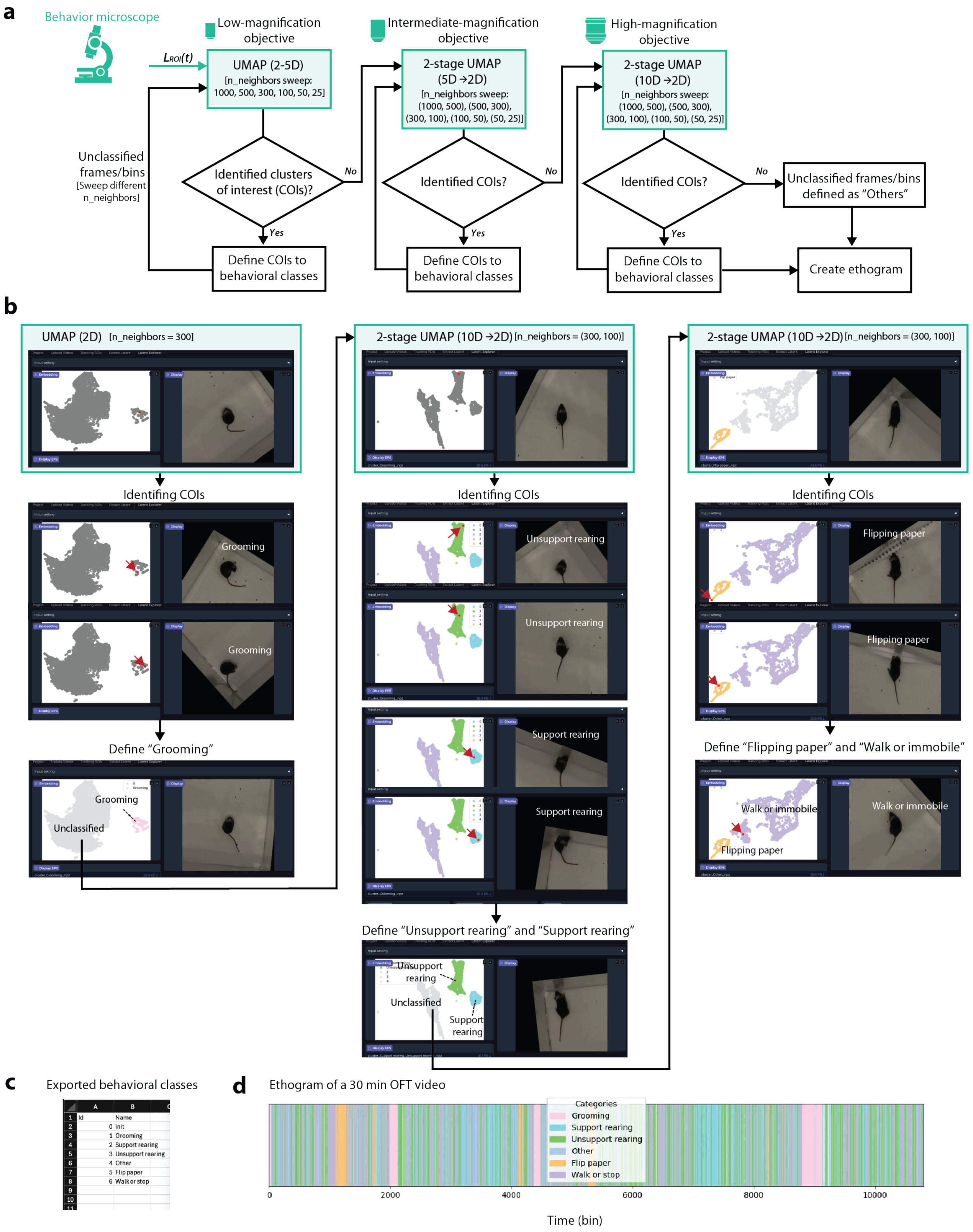


**Extended Data Figure 2. "Behavior Microscope": A hierarchical exploration framework for behavioral classification.**

**a**. Workflow diagram of the “Behavior Microscope”, illustrating the systematic progression through parameter spaces for behavioral cluster identification. **b**. Sequential screenshots demonstrating the interactive exploration process. The analysis begins with a "low-magnification objective" configuration to capture global behavioral structure. Through interactive visualization, users identify clusters of interest (COIs) by clicking on embedded points to examine corresponding video frames. In this example, the initial grooming cluster is identified and defined. When the same parameters fail to resolve additional COIs in the remaining data points, the system transitions to a "high-magnification objective" (two-stage *UMAP* with adjusted parameters), successfully revealing unsupported and supported rearing behaviors. Continued exploration with these parameters identifies a "flipping paper" behavior as an additional COI, while the remaining unclassified points comprise walking and immobile states. **c**. Representative output in CSV format containing cluster IDs and user-defined behavioral labels. **d**. Visualization of the resulting ethogram in PNG format, displaying the temporal sequence of classified behaviors across the recording session.

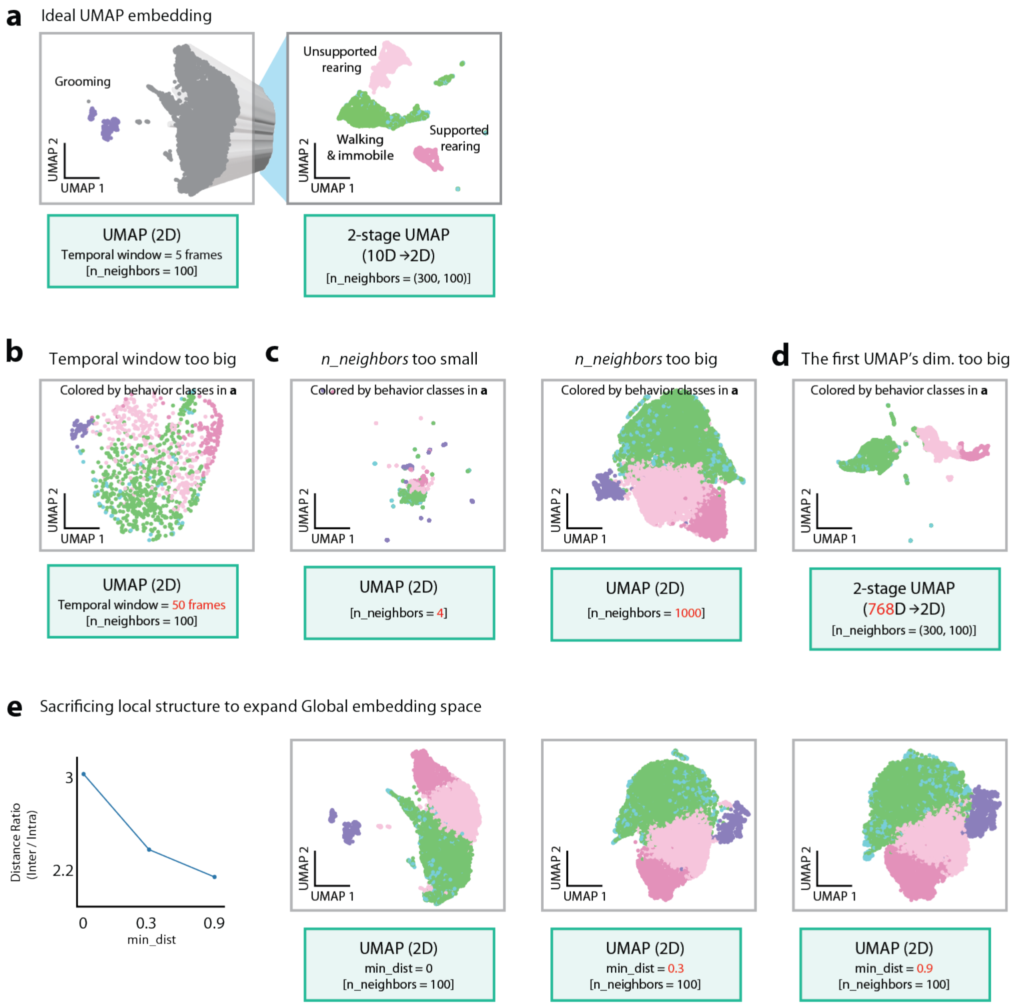


**Extended Data Figure 3. Guide to *UMAP* parameter tuning for behavioral clustering, illustrated with mouse OFT**

**a.** Example of well-separated behavioral clusters achieved with a set of *UMAP* parameters for mouse OFT data. **b.** Impact of an excessively large temporal window, causing all data to merge into a single, large undifferentiated cluster. **c.** Effects of the *n_neighbors* parameter: using a too low *n_neighbors* value results in over-fragmentation into many small clusters, while selecting a too large *n_neighbors* value obscures distinct groups by forming one large cluster. **d.** Influence of embedding dimension in the first *UMAP* step in a two-stage *UMAP* approach. A large first-stage *UMAP* embedding dimension, even if not perfectly optimized, distinguishes multiple behavioral clusters more effectively than a single-stage *UMAP* approach. **e.** Impact of the *min_dist* parameter on cluster separability. This panel illustrates how different *min_dist* values affect both the *UMAP* low-dimensional embeddings (visualizations) and the quantitative ratio of inter-cluster to intra-cluster distances. For the mouse OFT data shown, setting *min_dist* = 0 typically maximizes this distance ratio, resulting in the most visually distinct and well-separated behavioral clusters in the embedding.

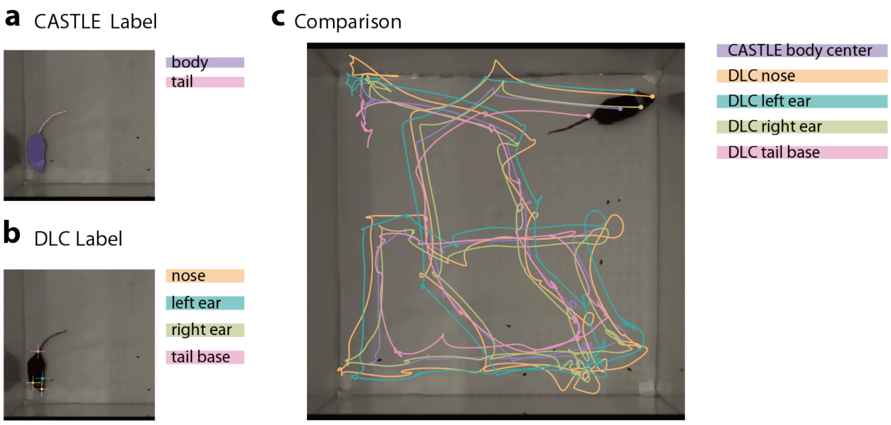


**Extended Data Figure 4. Comparison of tracking trajectories derived from *CASTLE* and *DLC*.**

**a.** Example of ROIs for the mouse body and tail, segmented and labeled using *CASTLE*. **b.** Key-points (nose, left ear, right ear, and tail base) tracked on the mouse using *DLC*. **c.** Overlay of movement trajectories. The trajectory defined by the center of the *CASTLE* body ROI (from **a**) is shown superimposed with a corresponding trajectory derived from the *DLC* key-points (from **b**), illustrating high consistency between the two tracking methods.

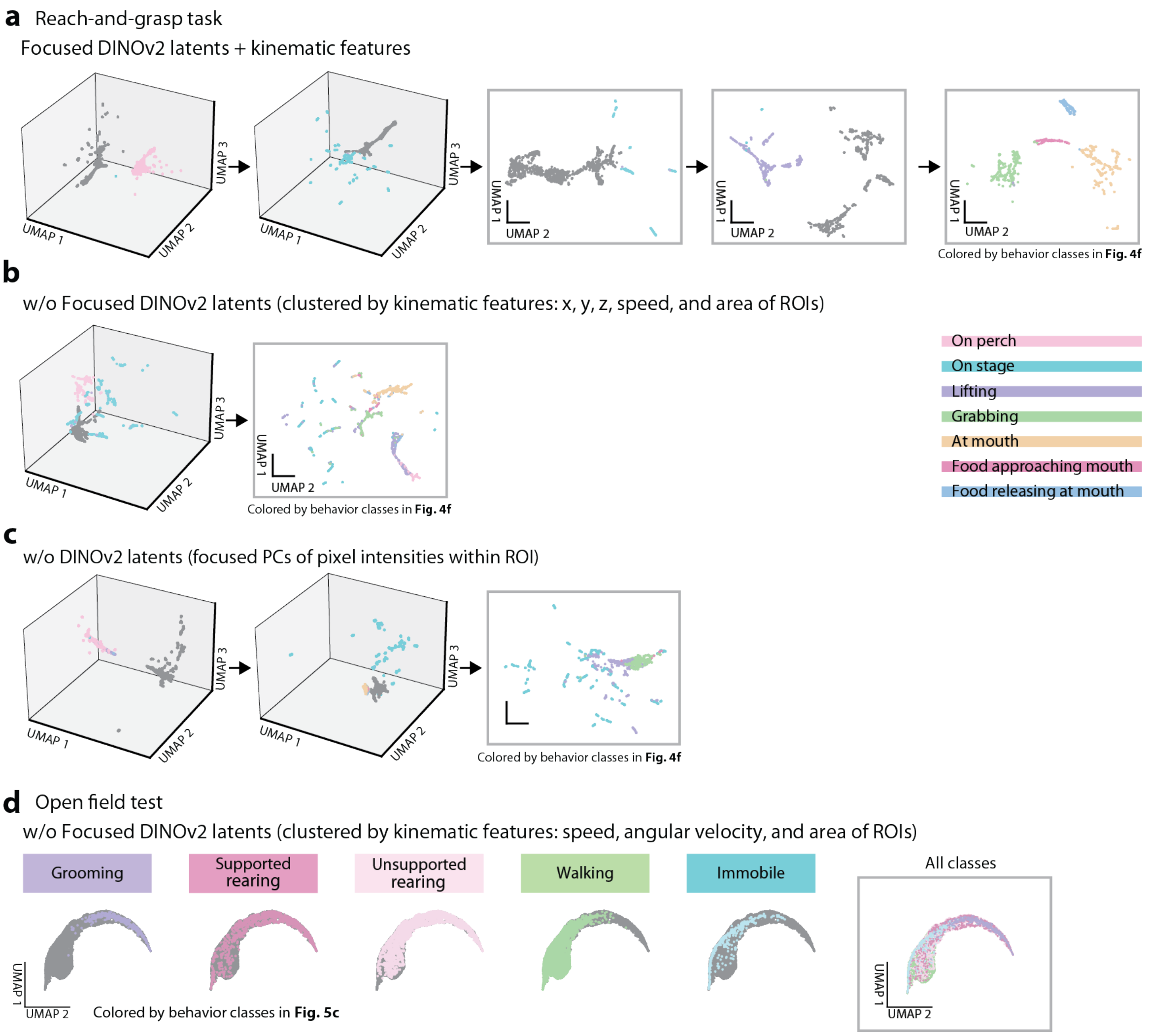


**Extended Data Figure 5. Ablation studies on feature engineering for behavioral analysis.**

**a.** *UMAP* embeddings from latent feature exploration. Using focused visual latent features combined with standardized kinematic features (3D position, ROI areas, speed). **b.** Necessity of ROI-based "focusing" for visual latent quality. The *UMAP* embedding was generated from *DINOv2* patch latent features averaged across the entire frame (without ROI-specific focusing). When points are colored according to behavioral classes identified using focused visual latent features, they show poor clustering and separation, emphasizing the importance of the focusing step. **c.** Contribution of *DINOv2*-based semantic features versus PCA on raw pixels. The *UMAP* embedding was generated from features derived by applying PCA directly to raw pixel values of cropped hand regions (front view: 60x60 px, side view: 100x100 px, each reduced to 768 dimensions). While some broad behaviors (e.g., "on perch," "on stage") can be distinguished, finer distinctions (e.g., between "on stage," "lifting," and "grabbing") are largely lost compared to using *DINOv2*-derived latent features, highlighting the value of semantic feature extraction for detailed behavioral analysis. **d.** *UMAP* embedding of mouse OFT behaviors using only kinematic features (speed, angular velocity, area). The significant overlap among all behavioral categories demonstrates that these kinematic features alone are insufficient for clear behavioral discrimination, underscoring the superior discriminative power of focused visual latent features.

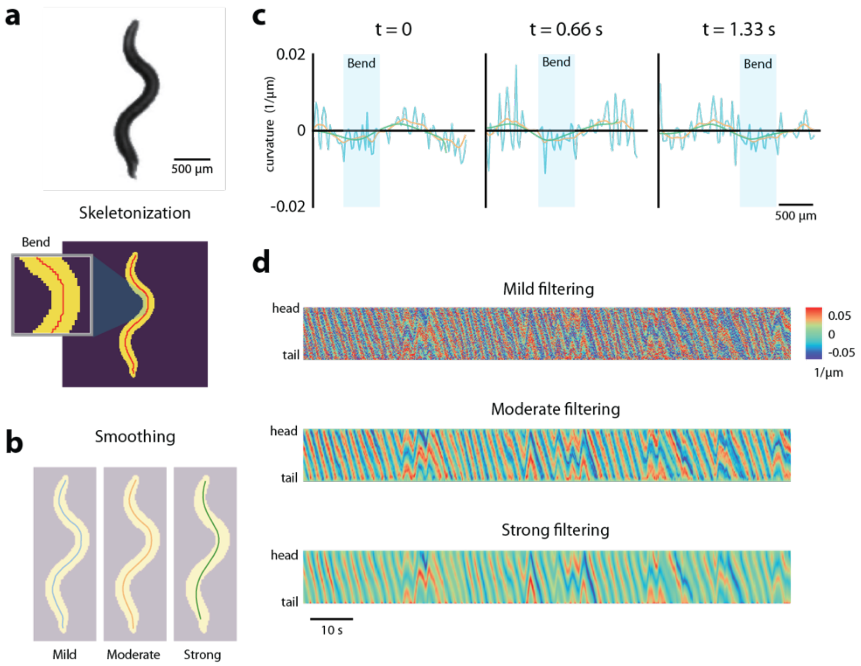


**Extended Data Figure 6. Optimization of *C. elegans* skeletonization and curvature analysis for behavioral validation**

**a.** Representative video frame of *C. elegans* with binary segmentation mask generated by applying Gaussian smoothing (5×5 kernel) and thresholding. The overlaid red line shows the raw thinned skeleton exhibiting characteristic step-like artifacts from pixel-based extraction algorithms. **b.** Skeleton refinement using low-pass filters of three distinct strengths: mild (sky-blue line, cutoff frequency 0.0159 $\mu m^{-1}$), moderate (orange line, 0.0053 $\mu m^{-1}$), and strong (green line, 0.00159 $\mu m^{-1}$) applied to x,y coordinates. **c.** Curvature profiles along the *C. elegans* body axis (~130 pixels) corresponding to the three filtering conditions. Moderate filtering demonstrates optimal balance between noise reduction and preservation of curvature dynamics. **d.** Temporal heatmaps of curvature dynamics along the worm body for mild, moderate, and strong filtering conditions over representative time windows.

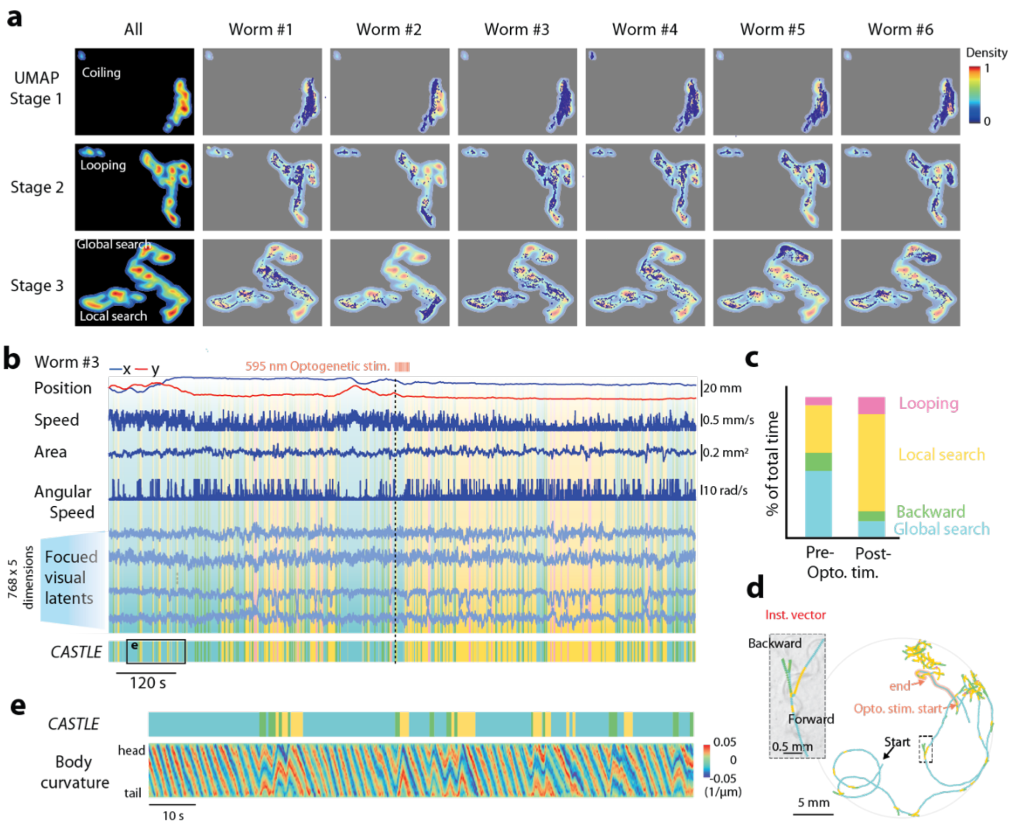


**Extended Data Figure 7. Variation of behavioral repertoires of *C. elegans* after optogenetic stimulation**

**a.** Inter-individual variability visualization showing behavioral state distributions projected onto three-stage *UMAP* embeddings for all worms (N = 6), revealing distinct behavioral selection repertoires across the worms. **b.** A additional example of temporal dynamics of kinematic parameters, focused visual latent features, and *CASTLE*-identified behavioral classes over the 20-minute recording from worm #3. **c**. Optogenetic stimulation effect on behavioral repertoire of worm #3. **d**. Optogenetic stimulation increases the “looping” and “local search” behavioral classes of worm #3. **e**. Example of body curvature dynamics tightly match *CASTLE* behavioral classifications.

### Supplementary Videos

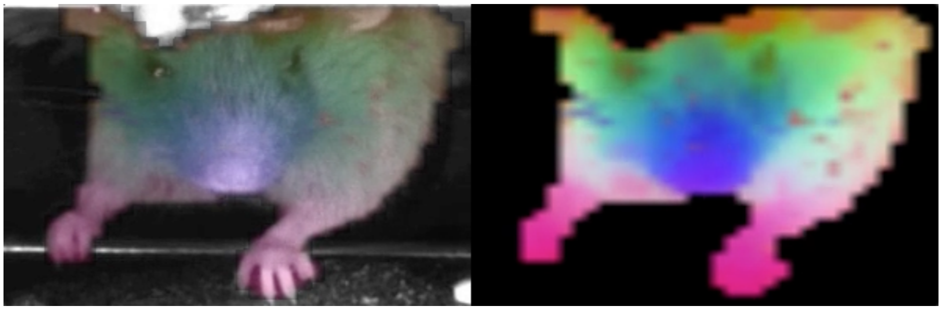

**Supplementary Video 1. Visualization of whole-body patch latent features using *PCA* in mouse reach-and-grasp task**

This video demonstrates how *DINOv2* encodes semantic visual information across the entire mouse body during a reach-and-grasp task. The patch latent features (37×37 patches, 768-dimensional each) are visualized using *PCA*, with the first three principal components mapped to RGB channels. The consistent color patterns across different frames reveal that *DINOv2* reliably identifies and encodes distinct anatomical regions, providing the foundation for subsequent ROI-focused feature extraction.

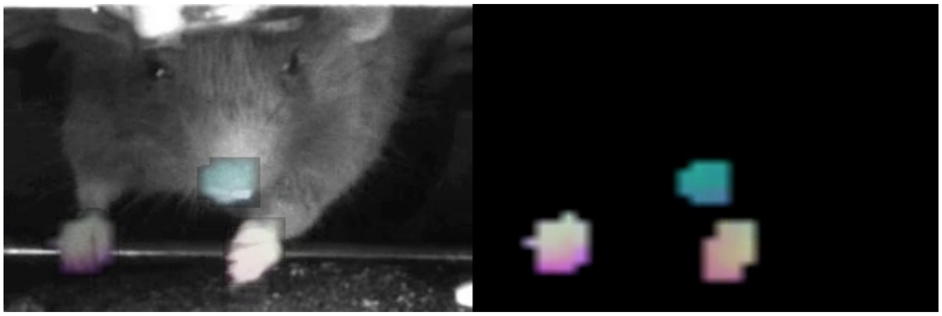

**Supplementary Video 2. ROI-focused patch latent feature visualization reveals distinct behavioral dynamics**

This video shows the PCA visualization of patch latent features specifically focused on three regions of interest (ROIs): left paw, right paw, and nose. The first three principal components are mapped to RGB channels, revealing how different ROIs exhibit distinct temporal dynamics during the reach-and-grasp sequence. Notably, the left paw (ROI1) shows dramatic color changes during reaching movements, while the right paw (ROI2) and nose (ROI3) maintain relatively stable representations, demonstrating the importance of ROI-specific feature extraction for capturing relevant behavioral information.

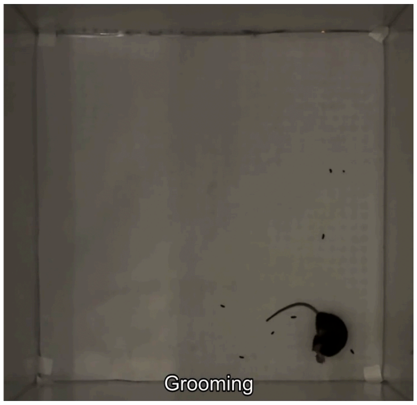


**Supplementary Video 3. Behavioral annotation visualization using *CASTLE*-generated subtitles**

This video demonstrates synchronized playback of *CASTLE*-generated behavioral class annotations with original mouse OFT footage. The subtitle track displays behavioral classifications (grooming, supported rearing, unsupported rearing, walking, and immobile states). This output from the "Behavior Microscope" workflow (**Extended Data Fig. 2**) enables validation of automated classifications against raw video data without requiring specialized software. The synchronized visualization facilitates verification of classification accuracy and identification of potential misclassifications at behavioral transition points.

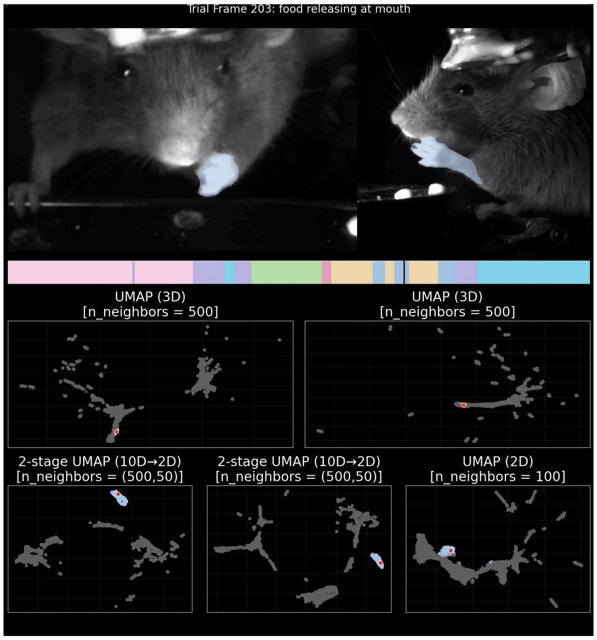


**Supplementary Video 4. Hierarchical *UMAP* exploration reveals fine-grained behavioral structure in mouse reach-and-grasp task.**

The video illustrates hierarchical *UMAP* embeddings of focused latent features at different levels of granularity during a mouse reach-and-grasp task. Although detailed reaching dynamics are not resolved in the initial stage 1 *UMAP*, hierarchical exploration reveals that this information is preserved and can be systematically uncovered, ultimately identifying stable behavioral classes including unexpected sub-actions not captured in the original annotations.

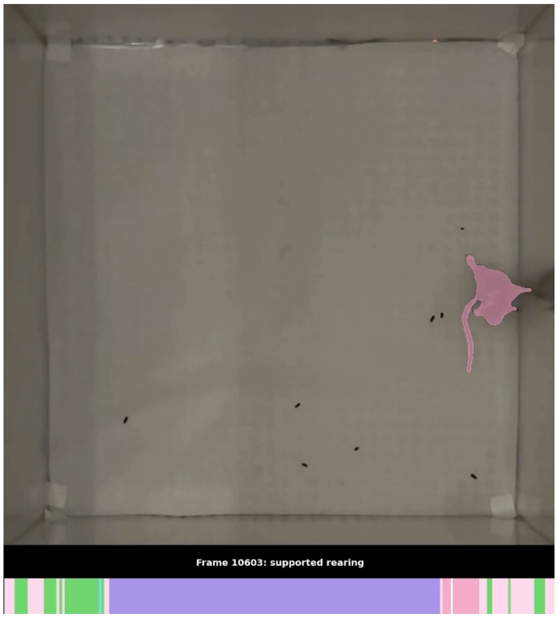


**Supplementary Video 5. Visualization of *CASTLE*-identified behavioral syllables with mouse open field test.**

This video show *CASTLE*'s behavioral classification results alongside the corresponding open field test video, enabling direct visual validation of the identified behavioral syllables. The original mouse behavior footage is shown with the temporal alignment of *CASTLE*-classified behavioral states (grooming, supported rearing, unsupported rearing, walking and immobile). The color-coded timeline indicates behavioral transitions throughout the recording, allowing readers to observe the correspondence between the automatically identified behavioral syllables and the actual mouse movements. This synchronized presentation demonstrates *CASTLE*'s accuracy in capturing subtle behavioral distinctions without requiring manual annotation.

### Method details

Here we describe the methodology for extracting focused visual latent features and the "Behavior Microscope" framework for interpreting these features. We then present the experimental protocols for each model organism, including the reach-and-grasp task and OFT in mice, as well as the behavioral assays conducted with *Drosophila* and *C. elegans*.

#### Core computational modules of *CASTLE*

<u>*Finding focused visual latent features*</u>

*CASTLE* integrates three core computational modules to enable comprehensive behavior analysis: (i) image prompting segmentation for initial ROI identification, (ii) video object segmentation for temporal tracking, and (iii) visual foundation models for extracting semantic features. This integration enables robust tracking and feature extraction across diverse experimental conditions.

To identify and track ROI from raw videos with dimensions H×W×C (height × width × channels), we first integrated image prompting segmentation and video object segmentation (VOS) techniques. For image segmentation, we employed Meta's *Segment Anything Model* (*SAM*), which identifies the ROI based on user-provided prompts (mouse clicks; **Fig. 1b**). For video object segmentation, we used *DeAOT* model[16]. which predicts the location of the ROI in subsequent frames based on the ROI defined in the first frame (**Fig. 1a and b**). Both *SAM* and *DeAOT* maintain the original video dimensions when outputting ROI prompts and ROI masks, ensuring no spatial information is lost during segmentation. Although Meta's recently released *SAM2*[49] has integrated both functionalities with improved performance, the combination of *SAM* and *DeAOT* in our study is already sufficient for effectively segmentation and tracking.

Next, to capture general visual semantic information, we adopted the visual foundation model *DINOv2*[17]. We use the base-size version of *DINOv2* with registers (approximately 86M parameters), which internally resizes all input images to 518×518 pixels regardless of their original dimensions—whether 300×200 or 500×500. With a patch size of 14×14 pixels, the model divides each frame into 37×37 patches (**Fig. 1a,c**). *DINOv2, using global image context, outputs a 768-dimensional feature vector for each patch, creating a* *feature tensor of dimensions 37×37×768 per frame. These patch-level features, referred to as patch latents, capture rich semantic information at each spatial location.* The base-size version of *DINOv2* is distilled from the 1.1-trillion-parameter giant-size *DINOv2* model and retains 98% of its performance, providing an optimal balance between computational efficiency and feature quality[17].

We provide the full mathematical descriptions of the pipeline in **Extended Data Fig. 1a**. In summary, to construct focused visual latents from the patch features, we developed a weighted averaging approach that preserves spatial correspondence between ROI masks and patch representations. First, the original ROI masks with dimensions H×W obtained from *DeAOT* are reshaped to match the 37×37 patch grid of *DINOv2*, generating the reshaped mask $\widetilde{ROI}(t)$. This reshaping process generates a weight matrix where each element $\widetilde{ROI}(t)_{ij}$ represents the proportion of the corresponding patch that overlaps with the ROI.

For each frame, the focused visual latent for an ROI is computed as:

$$L_{ROI}(t) = \frac{1}{N(t)} \sum_{i=1}^{37} \sum_{j=1}^{37} \widetilde{ROI}\,(t)_{ij} \, \cdot \, L(t)_{ij} \in R^{768} \tag{1}$$

Where the normalization factor $N(t)$ is defined as:

$$N(t) = \sum_{i=1}^{37} \sum_{j=1}^{37} \widetilde{ROI}\,(t)_{ij} \tag{2}$$

This weighted averaging ensures that patches fully within the ROI contribute more than those partially overlapping, resulting in a single 768-dimensional focused visual latent per ROI per frame. The normalization by $N(t)$ accounts for the variable number of patches covered by each ROI. This process transforms the spatial patch representation into a time-series dataset suitable for behavioral analysis (**Fig. 1e** and **Extended Data Fig. 1a**).

To visualize the rich information in the patch latent features, we performed PCA jointly on the patch latents corresponding to the mouse's left hand, right hand, and mouth regions. The values of the first three principal components were scaled to the 0–1 range and used as RGB values to visualize the corresponding ROI regions (**Fig 1c, d** and **Extended Data Figure 1d**). It is important to note that PCA was used here solely for visualization purposes. In the actual analysis pipeline, we preserved the full 768-dimensional features extracted by *DINOv2* to maintain maximum information content for downstream behavioral analysis.

### *Hierarchical clustering: "Behavior Microscope"*

For behavior analysis that requires temporal information, we employed an optional step, "temporal concatenation" (**Fig. 1e**). Specifically, we binned the high-dimensional focused latent vectors ($L_{ROI}(t)$ with dimensionality $D = 768$) from several consecutive frames, resulting in a high-dimensional feature matrix of size $b \times D$. Here, $b$ represents the bin size. Next, we applied *UMAP*[18] to embed the $L_{ROI}(t)$ into a low-dimensional space, where similar frames/bins naturally cluster together. To segment behaviors, we applied *DBSCAN* to the low-dimensional embedding, grouping nodes into distinct clusters, each representing a type of similar behavior. In *DBSCAN* algorithm, we employ a fixed epsilon parameter (eps = 1) as the default setting, which defines the maximum distance threshold for samples to be considered within the same neighborhood cluster.

However, projecting data directly to a two-dimensional space, a typical approach from previous work[6,39,40], may cause embedded signal to become overly aggregated. Minor differences between originally distinct nodes may be lost when dimensionality is too low, resulting in indistinguishable clusters. To address this issue, we propose a hierarchical, multi-stage *UMAP* strategy operationalized within the "Behavior Microscope" framework. This method allows for analysis at different conceptual magnifications: (i) Low magnification: To capture broad patterns and preserve the global data structure, we use *UMAP* to perform dimensionality reduction directly to low-dimensional space with a sweep of *n_neighbors* values from high to low (**Extended Data Fig. 2a**; see next session for detailed parameters). This is used to separate the most distinct behavioral classes first; (ii) High magnification: To resolve finer details within a cluster, we apply two-stage *UMAP*, which first embeds data to a 5- to 10-dimensional space, followed with a second *UMAP* to 2-dimensional space. This approach focuses on local relationships, revealing subtle sub-structures that were previously aggregated (**Extended Data Fig. 2b**). This two-stage process, moving from low to high magnification, gradually reveals and distinguishes subtler behavioral characteristics in a guided and interpretable manner.

The hierarchical approach for parameter selection, drawing an analogy to microscopy techniques with varying magnification objectives (**Extended Fig. 2a**). We set *min_dist* = 0 to minimize local structure constraints and prioritize global structure preservation (**Extended Data Fig. 3e**). The set of candidate values ($S$) for *n_neighbors* parameter, $S$ = {1000, 500, 300, 100, 50, 25}. We define the parameter spaces for two architectural configurations:

(i) Single-stage *UMAP*: The parameter space is

$$n_{\text{neighbor}} \in S \tag{3}$$

(ii) Two-stage *UMAP*: The parameter space is

$$n_{\text{neighbor}}^{(1)}, n_{\text{neighbor}}^{(2)} \in S, n_{\text{neighbor}}^{(1)} > n_{\text{neighbor}}^{(2)} \tag{4}$$

We typically start with a "low-magnification objective" approach to examine similarities and differences across all frames, thereby establishing a comprehensive understanding of the behavioral landscape. Subsequently, we systematically explore the *n_neighbors* parameter space in a coarse-to-fine manner. By progressively reducing the number of neighbor points, we obtain increasingly sparse topological structures in the low-dimensional embedding space. This process continues until clusters of interest (COI) emerge in the low-dimensional representation. If COI fail to materialize even at *n_neighbors* = 25, we transition to an "intermediate/high-magnification objective" approach utilizing the two-stage *UMAP* architecture (**Extended Data Fig. 2a, b**).

The rationale for this transition is that excessively small *n_neighbors* values compromise global structure preservation, causing similar behaviors to become dispersed across disparate regions of the low-dimensional embedding. The two-stage *UMAP* architecture addresses this limitation through a staged dimensionality reduction process, where the dimension parameter (*n_components* in *UMAP*) is progressively reduced: the 768-dimensional input is first reduced to an intermediate 5- or 10-dimensional representation, followed by a final reduction to 2 dimensions. Each stage serves a distinct functional role: the first stage employs larger *n_neighbors* values ($n_{\text{neighbor}}^{(1)}$) to preserve global structure and maintain proximity among similar behaviors, while the second stage utilizes smaller *n_neighbors* values ($n_{\text{neighbor}}^{(2)}$) to construct sparse topological structures that facilitate COI formation. The analysis pipeline generates two primary outputs: video subtitles and ethogram files (**Extended Data Fig. 2c, d** and **Supplementary Video 5**). Subtitles enable synchronized viewing of behavioral annotations with corresponding video footage for validation purposes. Ethogram data are provided in both CSV format for downstream computational analyses and PNG format for visual inspection of behavioral patterns. These standardized outputs ensure compatibility with existing analytical workflows (**Extended Data Fig. 2c, d**).

*Hierarchical clustering: parameter considerations*

When all nodes aggregate into a single large cluster in the low-dimensional space, making it impossible to identify the "most distinct clusters" (**Extended Data Fig. 3c**, *right panel*), we propose two alternative approaches. One involves reducing the number of similar bins (*i.e.*, neighbors) considered when constructing the *UMAP* graph, thereby creating a sparser graph. This approach helps separate nodes with inherently larger differences upon reconstruction. Another method is to employ two-stage dimensionality reduction with *UMAP*, gradually moving from higher to lower dimensions—for example, first reducing dimensions from 768 to 10, and then from 10 to 2 (**Extended Data Fig. 3a**, *right panel*). This helps effectively separate significantly different clusters. We observed that using too few neighbors could produce embeddings that lack global structural information. Hence, the two-stage *UMAP* strategy ensures a sufficient number of neighbors to capture global structure, while progressively reducing dimensionality to effectively distinguish local, subtle features. Although these strategies have demonstrated good results experimentally, the theoretical foundations behind them require further investigation.

**Mouse experiments**

C57BL/6J mice (8-12 weeks old) were used for the OFT. Mice were housed in groups of 4-5 per cage in a temperature-controlled environment (22 ± 1°C) with a 12-hour light/dark cycle. 3 weeks before behavioral assay, the mice were transfer to a room with reversed light/dark cycle. Food and water were available *ad libitum*. All procedures were conducted in accordance with the institutional guidelines and approved by the Institutional Animal Care and Use Committee (IACUC) at Academia Sinica.

**Mouse reach-and-grasp task**

*Experimental setup and data acquisition*

In this experiment, each trial lasted 30 seconds. Five seconds after the trial began, the mouse heard an auditory cue, after which a rotating plate delivered food in front of the mouse. The mouse then reached out to grab and eat the food. We analyzed the behaviors from 1 second before the cue to 5 seconds after the cue (total 6 seconds per trial). In most of the trials the mouse has put the food pallets into the mouth and the paw has returned to the rest stage, waiting for the next cue. Two high-speed IR cameras (Grasshopper 3, FLIR, USA) were used with 80 fps to collect data from both front and side views. The front view had a resolution of 512 x 344 pixels, where 6.58 pixels corresponded to 1 mm. The side view had a resolution of 768 x 502 pixels, where 11.16 pixels corresponded to 1 mm. Neural signals were recorded using a CMOS-MEA system [52] with a sampling rate of 32 kHz, and a total of 52 successful food-grasping trials were collected.

*Image processing*

Since our focus is on the mouse and its hand movements, we cropped the images from both perspectives to remove unnecessary portions, retaining only the regions that capture the full activity of the mouse's body. After cropping, the resolution of the front view is 300 x 200 pixels, and the side view is 500 x 500 pixels. Since *DINOv2* relies on semantic information to extract latent representations, we retained all mouse body parts within each frame, as we believe this approach helps the visual foundation model produce high-quality latent features.

*Neural signal processing*

We utilized Kilosort2[53] to perform spike sorting on the neural signals recorded 20 sessions (4 trials per session; total 80 trials) consisting a total of 2,300 seconds of neural signals. The spike sorting was performed by concatenating the signals from all sessions. The default parameters of Kilosort2 were used, with adjustments made to the threshold parameter ([8 3]) and the AUC parameter (0.8). We manually inspected the waveforms of each unit to eliminate false units, ultimately identifying 1,346 valid units. To align with the video's 80 fps, the spike train data of each unit were binned into 12.5 ms intervals, yielding spike count data at 80 Hz resolution based on their original spike times. 52 successful reach-and-grasp trials will be used in the following analysis.

*Behavior annotation and ROI tracking*

Two types of behavioral annotations were performed: keypoint tracking for precise hand trajectory and Region of Interest (ROI) segmentation for visual latent extraction.

For keypoint tracking, we used *DeepLabCut* (*DLC*) to label the mouse's left wrists in over 1,000 frames across both front-view and side-view videos to reconstruct hand trajectories (**Fig. 3a,b**). The *DLC* models were trained using default parameters. For ROI segmentation, we focused on identifying the mouse's left hand as the primary ROI across the 52 successful trials (each trial comprising 480 frames). Using our segmentation tools (*SAM* and *DeAOT*, as detailed previously), an average of 1.31 frames per trial (0.27% of total frames) needed to be manually annotated for the side-view videos. Notably, annotations from previously processed trials could be reused to aid tracking in subsequent trials. The need for more than one annotation per trial arose primarily from a common challenge in side-view videos: the proximity or occlusion of the left and right forelimbs. When the mouse lifted its left hand, the ROI sometimes split, with one part correctly tracking the lifted left hand and another erroneously drifting to the right hand (which was closer to the left hand's original position). This was resolved by providing corrective annotations on the affected frames, and these corrections were incorporated into the model's memory for subsequent trials (**Extended Data Fig. 1b**). For front-view videos, an average of 1.75 frames (0.35% of total frames) per trial required annotation. The primary issue here was the occasional disappearance of the hand ROI when the mouse brought its paw to its mouth, as the model sometimes misidentified the hand as part of the mouth. The centroid of these segmented ROIs was also used to construct movement trajectories (**Fig. 3a,b**).

We observed that during periods when the mouse was stationary, the *DLC*-generated keypoint labels occasionally exhibited minor jitter, an issue also noted in other studies such as *Keypoint-MoSeq*[38] (**Fig. 3c**). To compare tracking stability, we analyzed the speed variability (jitter) during periods when the mouse's hand was physically stationary. From the reach-and-grasp task, we calculated the 2D speed (using x and y coordinates from the side view) for the time interval prior to the onset of each reach. We identified and selected the 44 out of 52 trials where the hand was confirmed to be stationary during this pre-reach phase (**Fig. 3d**). For each of these 44 trials, we computed the standard deviation of the speed trace derived from each tracking method (e.g., *CASTLE* vs. *DeepLabCut*) as a quantitative measure of variability. A paired Wilcoxon signed-rank test was then used to determine if there was a statistically significant difference between the standard deviations of the paired speed measurements.

*Decoding hand position from neural activity*

A recurrent neural network (RNN) was employed to decode left-hand position from neural activity. The network architecture consisted of two Gated Recurrent Unit (GRU) layers (embedding dimension = 64) followed by two fully connected (FC) layers. The model's input was the binned spike counts from 1,346 validated units over a temporal window of the 20 preceding frames. The output was the predicted X and Y coordinates of the left hand, trained against ground-truth trajectories derived separately from both *DLC* and *CASTLE* (**Fig. 3e**).

For training, the dataset of 52 successful trials was split into a training set (trials 1–36), a validation set (trials 37–42), and a test set (trials 43–52). The model was trained for 100 epochs with a batch size of 64, using the Mean Squared Error (MSE) loss function and the Adam optimizer with a learning rate of $1 \times 10^{-4}$. To assess the stability of the decoding performance, the entire training and evaluation process was repeated 10 times with independent initializations. The resulting decoding performance (measured by Coefficient of determination, $R^2$) and loss traces were then analyzed to evaluate the downstream impact of tracking jitter from the different ground-truth sources.

*Latent exploration and behavioral classifications*

First, we defined five target behaviors—on perch, lifting, grabbing, at mouth, and on stage—before beginning latent exploration. Each view (front and side) produced a 768-dimensional feature vector, and we concatenated these to form a 1,536-dimensional feature representation for each frame (**Fig. 4a**).

We then adopted a hierarchical clustering strategy using *CASTLE*'s Latent Explorer to systematically investigate behavioral classes from the focused visual latent features (**Fig. 4b**). In the first stage, *UMAP* (n_neighbors = 500, *n_components* = 5) was applied for dimensionality reduction to reveal the overall behavioral structure. This initial step successfully isolated a cluster. By observing the corresponding video frames for the data points within this cluster, we identified the behavior and named it the "On perch" cluster (pink), which was then separated from all other behaviors for subsequent analysis. In the second stage, we reused the same *UMAP* parameters to further isolate “On stage.” (blue) For the third stage, we adjusted *UMAP* to (*n_neighbors* = 100, *n_components* = 2) to enhance local resolution. By constructing a sparser *UMAP* graph, we distinguished “On stage” frames that had previously clustered with other behaviors.

In the fourth stage, we ran a two-layer *UMAP*: layer 1 (*n_neighbors* = 500, n_components = 10) to preserve global structure, and layer 2 (*n_neighbors* = 50, *n_components* = 2) to refine local boundaries. This allowed us to separate “Lifting” (purple) from the remaining behaviors. Observing residual structure in the embedding, we then removed the “Lifting” points and reapplied the same parameters. At this point, we clearly distinguished “grabbing” (green) and “at mouth” (yellow), with “lifting” frames positioned between these two clusters. In fact, the “lifting” behavior identified in Stage 5 can be subdivided into two actions: “food approaching mouth” and “food releasing at mouth” (**Fig. 4f–g**).

*<u>Validation of behavioral classifications & decoding behavioral classes</u>*

We used two methods to validate *CASTLE's* classification performance: (1) comparing *CASTLE* labels to manual labels using similarity metrics, and (2) assessing how well neural activity can predict both manual and *CASTLE* labels, thereby evaluating whether they carry correct information content.

First, we calculated the weighted F1 score between manual labels and *CASTLE* labels, obtaining a score of 0.90 (**Fig. 4d**). Next, to predict behavior from neural activity, we constructed a classifier with a two-layer GRU followed by a two-layer fully connected (FC) network. This classifier takes binned spike counts from the past 19 frames and now total 20 frames as input and outputs the likelihood for each of the five behavior categories at each time point. The embedding dimension was set to 1,024, and we trained for 60 epochs with a batch size of 256. Of these trials, 36 were used for training with an *Adam* optimizer (learning rate = $1 \times 10^{-5}$), 6 for validation (to prevent overfitting), and 10 for testing. We trained and evaluated the classifier separately on two label sets: manual labels and *CASTLE* labels. For each label set, predictions were generated 10 times, and we computed the F1 scores for all five behavior categories. The resulting F1 scores from these 10 runs were then statistically compared between the manual and *CASTLE* label sets for each behavior class using the Wilcoxon rank-sum test (**Fig. 4e**).

**Mouse OFT**

*Experimental setup and data acquisition*

The open field arena consisted of a square Plexiglas box (40 cm × 40 cm × 40 cm) with opaque walls. The arena was cleaned with 70% ethanol between trials to remove olfactory cues before performing any test. Each mouse was gently placed in the center of the arena at the start of the test. Videos were recorded at 1080p 30 fps using an iPhone 15 Pro mounted directly above the box. The first 5 minutes after placement were excluded from analysis. Control mouse was recorded for 2 hours, while PD and mild model mice were recorded for 1 hour. For all three animals, behavior was analyzed within a 30–60-minute window. The room was kept quiet with dim light to minimize external disturbances, and tests were conducted during the dark phase of the circadian cycle to ensure the mice were in their active state.

*Behavior annotation and ROI tracking*

Videos was cropped to 720 × 720 pixels to cover the open field arena only. For *DLC*, we annotated 220 frames, marking the mouse’s nose, left and right ears, and tail base (**Extended Data Fig. 3 b, c**). In *CASTLE*, two ROIs—the body and the tail—were tracked for aligning each frame. The first frame and two additional random frames were annotated. In the case of ROI losing by hidden tails or motion blur, extra frames (11 frames in open field tests) were annotated to complete the tail tracking. Using the Orientation Neutralization method, extra annotations is not required. To make the body is always upward, each video frame was aligned by rotating the frame. Frames were cropped to

420 × 420 pixels for extracting focused visual latent features, effectively removing any orientation information.

#### *Latent exploration and behavioral classifications*

Initially, 4 targeting behaviors were defined: walking, rearing, grooming, and immobility. An excessively large temporal window led to behavioral clusters becoming overly dense (**Extended Data Fig. 3b**). To improve cluster separation, the temporal window was set to 5, that consecutive frames were concatenated into a single temporal bin.

In the first step of "Behavior Microscope" analysis, we applied *UMAP* (*n_neighbors* = 100, *n_components* = 2, temporal window = 5) and successfully separated grooming from the other behaviors. In the second step, we ran a two-stage *UMAP*: Stage 1 (*n_neighbors* = 300, *n_components* = 10) to preserve global structure and Stage 2 (*n_neighbors* = 100, *n_components* = 2) to refine local boundaries. In this step, two behavioral subtypes—unsupported rearing and supported rearing—were split. To distinguish behaviors that are difficult to separate based solely on focused visual latent features (e.g., walking vs. immobility), a speed threshold[25] of 4 mm/s was set, yielding five distinct categories in total (**Fig. 5c**). Same analysis was applied to both mild and severe PD mice. The 10-minute OFT trajectories (**Fig. 5e**) and total behavior durations (**Fig. 5f**) were used to assess and compare behavioral patterns across different severities.

#### *6-OHDA mouse model of Parkinson's disease (PD)*

Two adult mice (12-20 weeks) were used for hemi-parkinsonian induction. Mice were first anesthetized by 5% isoflurane for induction. After placing the mice on the stereotaxic frame, 1.5% isoflurane was given to keep the mice be anesthetized. The fur was removed, and the skin was sterilized by iodine solution and 75% alcohol. After incision of the skin over the skull, a hole relative to bregma: AP -1.2 mm, ML: 1.1 mm, was drilled for creating access to medial forebrain bundle (MFB). Different severity of PD-mouse was created by injection of different 6-OHDA concentration through glass needles at depth: DV 5.0 mm relative to cortical surface. For the weak PD model, 0.2 µl final concentration (12.5 mg/ml) was injected with 6-OHDA dissolved in 0.2 % ascorbic acid, while severe PD model was induced by injection of 1 µl final concentration (3 mg/mL) with 6-OHDA dissolved in 1%

ascorbic acid. To minimize the toxicity of 6-OHDA to norepinephrinergic neurons, desipramine (25 mg/kg) was injected (*i.p.*) 30 min prior to the surgeries.

*Neutralization of the orientation effect of visual latent encoding (Orientation Neutralization)*

Due to the inherent limitations of alignment-based approaches, certain experimental scenarios present significant challenges for maintaining consistent body orientation. Specifically, cases where the head and tail of *C. elegans* become physically connected prevent reliable orientation maintenance through naive alignment methods for eliminating orientation-dependent information. To address this methodological constraint, we propose a novel approach termed Orientation Neutralization.

Initially, we conducted a systematic investigation to confirm the presence of orientation effects within the latent feature representations. We rotated a single frame of mouse locomotion through a complete 360-degree cycle, subsequently analyzing the variance across each latent dimension to quantify orientation-dependent effects (**Fig. 6a**). Additionally, we employed PCA to comprehensively evaluate the collective influence of all dimensions within the focused visual latent feature space (**Fig. 6b**).

Subsequently, we performed a comparative analysis of each preprocessing step involved in orientation effect removal, examining their respective impacts on latent feature representations and latent exploration. We first plotted our baseline performance using the previously validated method consisting of Focus combined with Alignment (**Fig. 6c-d**). We then implemented a two-stage latent exploration approach (employing identical parameters as specified in the *OFT Latent Exploration and Behavioral Classifications* chapter) to demonstrate the effects of each preprocessing step on latent exploration outcomes. This analysis included raw OFT frames (**Fig. 6e-f**), Focus preprocessing (**Fig. 6g-h**), and Orientation Neutralization (**Fig. 6i-j**).

The proposed orientation neutralization approach involves systematic rotation of the cropped image around the centroid of the body ROI, extracting focused visual latent representations at 15-degree intervals. These multiple representations are subsequently averaged to neutralize orientation-dependent influences.

For performance evaluation, we utilized behavioral sequence classifications derived from the alignment method as our baseline reference standard. To assess the efficacy of

Orientation Neutralization, we examined whether the generated classifications matched the alignment-based labels from identical or temporally adjacent time bins. Matching predictions were classified as correct, while non-matching predictions retained their original classifications. Using this evaluation framework, we computed weighted F1 scores (**Fig. 6k**).

Through comprehensive analysis of latent exploration results and behavioral class timelines, we observed that classification errors frequently occurred during behavioral transition periods. Consequently, we calculated F1 scores incorporating temporal tolerance for transition-related errors. For each time point classified by the alignment method, we searched for matching orientation effect classification results within ±T time bins, designating matches as correct classifications. We generated curves depicting F1 score performance across varying time bin tolerance levels (**Fig. 6l**) and visualized the confusion matrix under ±2 bin (±330 milliseconds) tolerance conditions (**Fig. 6m**).

*Ablation studies for feature evaluation*

To comprehensively evaluate the contributions of different components within our feature engineering pipeline, we conducted a series of analytical comparisons and ablation studies (**Extended Data Fig. 5**).

First, we explored the performance of integrating kinematic data with our focused visual latent features (**Extended Data Fig. 5a**). This involved standardizing a set of kinematic features—comprising 3D coordinates (x, y, z) of the region of interest (ROI), ROI area derived from both front and side camera views (area of front view, area of sideview), and movement speed—using their respective standard deviations. These standardized kinematic features were then concatenated with the *DINOv2*-derived focused visual latent features. The resulting combined feature vectors were subsequently used as input for the downstream latent exploration process, involving *UMAP* embedding and clustering, to observe their influence on behavioral cluster formation.

Subsequently, we investigated the necessity of the 'focus' aspect within the "focused visual latent features" (**Extended Data Fig. 5b**). To achieve this, an alternative feature set was generated by averaging *DINOv2* patch latent features from the entire video frame for each frame, thereby creating a global, non-focused visual feature vector. These global frame latent features were then processed through the identical latent exploration pipeline.

The quality of the emergent behavioral clusters from this approach was assessed by applying class labels that were originally identified using the ROI-focused visual latent features.

Furthermore, the significance of employing a semantic feature extractor like *DINOv2* was examined by comparing its output against a simpler pixel-based representation (**Extended Data Fig. 5c**). For this comparison, the ROI corresponding to the left hand was cropped from original video frames based on its center (60x60 pixels for the front view, 100x100 pixels for the side view). Principal Component Analysis (PCA) was then applied directly to the raw pixel values of these cropped image patches from each view separately, reducing their dimensionality to 768 to match the *DINOv2* feature dimension. The concatenated PCA-derived features from both views were subsequently used for latent exploration and behavioral clustering.

Finally, to ascertain the discriminative power of purely kinematic information, particularly for mouse OFT behaviors, we utilized a feature set consisting solely of speed, angular velocity, and body area (**Extended Data Fig. 5d**). These kinematic features alone were input into the latent exploration and clustering pipeline, and their ability to differentiate distinct OFT behavioral categories was evaluated.

### Behavioral analysis in *Drosophila*

#### Animals and experimental setup

*Drosophila melanogaster* was reared under standard conditions on cornmeal food at 23$^{o}$C under 12h/12h light/dark cycle and 60% relative humidity. All flies used for tracking were 5-6day old mated Canton-S females. Each fly was gently blown into a circular chamber with diameter of 5cm and approximate height of 1.8mm. The chamber featured a transparent ceiling and walls placed above an infrared (IR) bottom light panel. Individual flies were then positioned in front of a laptop (M2 MacBook air, 500nits brightness) with a screen inclined at 57.5$^{o}$ angle. The setup was placed in a room illuminated by redlight. After 20second of acclimation period,

a white screen of 1200(h) x 2000(w) resolution is displayed. Fly behavior was recorded for 3minutes using a Blackfly S BFS-U3-13Y3M camera at 60 fps.

#### *Image processing and ROI tracking*

Due to a circular bright band present in the center of our recording arena (**Fig. 7a**), we first performed background subtraction to ensure that this artifact did not interfere with the semantic feature extraction by the focused visual latent model. We randomly selected 10% of the frames from each video and calculated the median pixel intensity value for each pixel position to generate a static background model. Given that the flies appear as dark objects, the pixel values were inverted (e.g., 0 becomes 255). This step aligns the data representation with the intuitive concept that higher numerical values correspond to the signal of interest (the fly), while lower values represent the background. The median background was then subtracted from these inverted frames. Since subtraction could result in negative values, the pixel intensities were clipped to a range of 30-255. Finally, the pixel values were inverted again, returning the fly pixels to lower intensity values.

Following background subtraction, we annotated the first frame and two additional randomly selected frames per video for Region of Interest (ROI) tracking.

#### *Focused visual latent extraction and behavioral analysis*

To extract focused visual latent features while mitigating orientation-dependent biases, we employed an Orientation Neutralization strategy. This involved rotating the image around the ROI's center and cropping it to 210x210 pixels for each of the 24 rotational steps (15° increments) before latent extraction with *DINOv2*, then averaging these latent features.

A single-stage *UMAP* (parameters: *n_neighbors* = 25, *n_components* = 2, *temporal window* = 10) was sufficient to explore and identify distinct behavioral clusters (**Fig. 7c**). To validate these clusters and their corresponding behavioral interpretations, we visualized the standard deviation of pixel changes within a 15-frame interval, mapping this activity measure onto the corresponding video frames (**Fig. 7e**). Finally, we analyzed the probability distribution of behaviors on the *UMAP* embedding to observe behavioral preferences across the three individual flies (**Fig. 7f**).

### **Behavioral analysis in *C. elegans***

*Animals and optogenetic stimulation*

*C. elegans* at age day 1, strain: NTU0032, with genotype *him-5(e1490)/V; chcEx011[Ppkd-2::Chrimson; Posm-6::GFP]* was used for optogenetic experiments. This strain carries the extrachromosomal array *chcEx011*, in which the red-light-activated ion channel Chrimson is expressed under the *pkd-2* promoter in ciliated sensory neurons, with GFP driven by the *osm-6* promoter serving as a co-injection maker. The *him-5(e1490)* mutation provides a genetic background that increases the proportion of male worms for experimental convenience. L4 larvae were collected one day prior to experiments and cultured on nematode growth medium (NGM) plates containing *Escherichia coli* strain OP50 and 2 mM all-trans-retinal (ATR, Sigma-Aldrich, CAS number: 116-31-4). On the experimental day, healthy adult worms expressing GFP fluorescence were selected for behavioral assays using the *WormLab* system (MBF Bioscience). Individual animals were placed on NGM culture plates spotted with 20 µl of *E. coli* strain OP50 and recorded using a Basler acA2440 camera at 2448 × 2048-pixels resolution with a spatial resolution of 18.87 µm/pixel. Optogenetic stimulation was performed using 595 nm orange light at an intensity of 0.4283 $mW/mm^2$. Each 20-minute video recording included a light stimulation protocol that began at 9 minutes and 45 seconds. The stimulation paradigm consisted of 500 ms light pulses followed by 1-second intervals, creating 1.5-second complete stimulation cycles. A total of six videos were recorded. A total of 20 light stimulations were delivered, with the entire stimulation phase concluding at 10 minutes and 15 seconds. Given that behavioral changes were anticipated, videos were recorded to analyze behaviors before and after optogenetic stimulation.

*Image processing and ROI tracking*

To focus specifically on the postural dynamics of *C. elegans* locomotion, we first performed ROI tracking. The initial frame and two additional randomly selected frames were annotated to initialize the ROI tracking. After obtaining the ROI for each frame, a masking procedure was applied to create a new video sequence containing only the C. elegans' posture within the ROI, effectively removing all background information. This masking approach was chosen over the median background subtraction method (used for Drosophila) because *C. elegans* leave persistent trails during locomotion that cannot be adequately removed by a static background model.

*<u>Focused visual latent extraction and behavioral clustering</u>*

To extract focused visual latent features from background-removed worm images, we implemented an orientation neutralization strategy. This approach involved rotating each image around the region of interest (ROI) center through 24 discrete angular steps, cropping to 140×140 pixels at each rotation, extracting latent features, and computing their average focused visual latent features (**Fig. 8a**).

Subsequently, we employed a hierarchical clustering strategy to systematically categorize behaviors from the extracted orientation-neutralized focused visual latent features. To evaluate the discriminative power of *CASTLE*, we performed a comparative analysis with traditional kinematic parameters. In the case of worm, we calculated angular speed, defined as the rate of angular change between two consecutive line segments: the first connecting positions at time points t-1 and t, and the second connecting positions at time points t and t+1. This kinematic measure, along with speed and ROI area, provided a baseline locomotor dynamic for comparison with our focused visual latent representations (**Fig. 8b**). The analysis proceeded through multiple stages:

In the initial stage, we applied *UMAP* (*n_neighbors* = 30, *n_components* = 2, *temporal window* = 5), to reduce dimensionality and reveal the underlying behavioral structure. This preliminary analysis successfully isolated a distinct cluster, which, through manual inspection of corresponding video frames, was identified and designated as the "coiling" behavioral state (red). This cluster was subsequently excluded from further analysis. The second stage employed a two-layer *UMAP* architecture to achieve refined behavioral segregation. Layer 1 (*n_neighbors* = 300, *n_components* = 5) preserved global structural relationships, while Layer 2 (*n_neighbors* = 100, *n_components* = 2) enhanced local boundary definition. This approach enabled the identification and separation of the "looping" behavioral cluster (pink) from the remaining frames. Subsequently, applying identical parameters, we successfully distinguished the "local search" cluster (**Fig. 8c**).

Then, distinguishing between backward and forward global search presented particular challenges, as backward movements frequently manifest between global search episodes and often exhibit S-shaped trajectories similar to forward global search patterns.

This morphological similarity renders discrimination based solely on focused visual latent features insufficient. To address this limitation, we incorporated positional trajectory data, which exhibits characteristic fluctuations due to the worm's lateral undulations. To attenuate high-frequency noise, we applied a second-order Butterworth low-pass filter with a cutoff frequency of 0.5 Hz to the positional data. For directional classification, we defined three positional coordinates at each time point: the historical location $P_{ref}$, the current position ($P_t$), and the immediate past position ($P_{t-1}$). The historical vector was computed using the historical location and the current position, while the instantaneous vector was derived from the immediate past and current positions. Directionality was determined based on the dot product of these two vectors. Points already classified into the coiling, local search or looping categories were ignored, whereas unclassified points were assigned either a forward or backward label. Initially, the $P_{ref}$ was set to the current position $P_t$. As the worm moved, the historical location was updated whenever the displacement from the current position exceeded a predefined threshold D, until the distance between the historical location and current position became less than D. The historical location was reset only during initialization or upon occurrence of looping behavior. In our analysis, the displacement threshold D was set at 2.5 mm. See **Algorithm 1** for implementation details (**Fig. 8d**).

**Algorithm 1** Compute Directional Dot Products

threshold $\leftarrow D$
index_ref $\leftarrow 0$
pos_ref $\leftarrow$ pos[0]
dot_values $\leftarrow$ empty list
**for** $i \leftarrow 1$ to length(pos) $- 1$ **do**
  **if** data[$i$] = CLUSTER_ID_LOOPING **then**
    index_ref $\leftarrow i$
    pos_ref $\leftarrow$ pos[$i$]
    **continue**
  **end if**
  **while** dist(pos_ref, pos[$i$]) $\geq$ threshold **do**
    index_ref $\leftarrow$ index_ref + 1
    pos_ref $\leftarrow$ pos[index_ref]
  **end while**
  $\delta_{\text{inst}} \leftarrow \text{pos}[i] - \text{pos}[i-1]$
  $\delta_{\text{hist}} \leftarrow \text{pos}[i] - \text{pos\_ref}$
  $\mathbf{u}_{\text{inst}} \leftarrow \frac{\delta_{\text{inst}}}{\|\delta_{\text{inst}}\|}$
  $\mathbf{u}_{\text{hist}} \leftarrow \frac{\delta_{\text{hist}}}{\|\delta_{\text{hist}}\|}$
  append(dots_values, $\mathbf{u}_{\text{short}} \cdot \mathbf{u}_{\text{long}}$)
**end for**

The behavior selections (**Fig. 8g**) served as the basis for quantitative comparison of locomotor behaviors before and after optogenetic stimulation, with error bars indicating the standard error of the mean (SEM).

To characterize individual variability in behavioral expression, we projected data points from each animal onto the three-stage *UMAP* embeddings, thereby visualizing the distribution of inter-individual behavioral differences (**Extended Data Fig. 7a**). Representative examples are presented for worm #3 (**Extended Data Fig. 7b-e**) and worm #4 (**Fig. 8b, f, h**), displaying their respective behavioral syllables, trajectories, and body curvature profiles. These examples illustrate animals exhibiting distinct behavioral selection repertoires, specifically those with and without coiling behavior, respectively.

*Behavioral class validation using curvature analysis*

To validate the behavioral classes identified by *CASTLE*, we performed a detailed curvature analysis of the worm's posture (**Fig. 8h**). First, all frames were initially smoothed using a Gaussian blur (5×5 kernel) to reduce noise before segmentation via Otsu's thresholding. The skeleton was extracted using the *cv2.ximgproc.thinning()* function from the OpenCV library. Raw skeletons produced by this method often exhibit step-like artifacts due to the pixel-based algorithm, which introduces high-frequency noise into curvature calculations (visualized in **Extended Data Fig. 6a**). To mitigate this, we applied a low-pass filter to the x- and y-coordinates of the extracted skeleton. We explored three filter strengths to find an optimal balance between noise reduction and signal preservation: mild (cutoff frequency: 0.0159 $\mu m^{-1}$), moderate (0.0053 $\mu m^{-1}$), and strong (0.00159 $\mu m^{-1}$). Upon evaluating the resulting curvature profiles and temporal dynamics (**Extended Data Fig. 6b-d**), the moderate filter was selected as it provided the optimal balance, effectively reducing noise while preserving genuine curvature dynamics. This optimized curvature information was then used as a quantitative ground truth to validate the *CASTLE*-derived behavioral classes.

**Statistical analysis**

To examine the effects of stimulation (Pre vs. Post, N=6) and behavior type on the proportion of time spent in each behavioral state, we employed a two-way repeated-measures nonparametric analysis using the *aligned rank transform (ART)* procedure, implemented in the *ARTool* package in *R*. This approach allows for the analysis of main effects and interactions in factorial designs without assuming normality, by aligning and ranking the data separately for each effect prior to *ANOVA*. The within-subject factors were stim (two levels: Pre, Post) and behavior (five levels: Coiling, Looping, Global Search, Local Search, Reverse), with subject as the repeated-measures blocking factor. Following the *ART ANOVA*, we conducted post-hoc pairwise comparisons using *emmeans* with *Tukey*'s adjustment for multiple testing. Specifically, simple main effects of stimulation were tested within each behavior type, and pairwise comparisons among behaviors were performed within each stimulation condition (**Fig. 8g**). Statistical significance was set at $p < 0.05$ for all analyses. All statistical tests were two-tailed.

**Computational requirements and performance**

*Hardware specifications*

For optimal performance of the *CASTLE* pipeline, we recommend a minimum hardware configuration equipped with an NVIDIA RTX 3070 GPU (8 GB VRAM). The analyses presented in this study were conducted using an NVIDIA RTX 4090 GPU (24 GB VRAM) and Intel Core i5-12600KF processor.

<u>*Computational time requirements*</u>

To illustrate the computational demands of each processing stage, we provide timing benchmarks using a representative video of mouse open-field test (30 minutes duration, 30 fps, 1000×1000 pixels, total 54,000 frame) as an example. In summary, ~90 minutes are required to obtain the final behavioral classification results. The step-by-step break down of the time benchmarks are as following:

1. Interactive ROI segmentation: Near real-time performance with mask generation completed within seconds per frame, enabling efficient annotation of reference frames. (< 1 minute)
2. Video object segmentation: Full video tracking using *DeAOT* (a *ResNet50* backbone) required approximately 41.5 minutes to propagate ROI masks across all 54,000 frames while maintaining temporal consistency. (41.5 minutes)
3. Focused visual latent extraction: Processing time varied significantly based on the preprocessing strategy employed. For alignment-based method, it takes ~21.5 minutes, and for orientational neutralization (24-step rotation), it takes 24 times of alignment-based method.
4. Behavioral clustering: GPU-accelerated *UMAP* dimensionality reduction completed within seconds per embedding, enabling rapid iterative exploration of clustering parameters. (10 – 30 minutes)

To benchmark *CASTLE*'s computational efficiency, we compared it against the established *DLC* + *Keypoint-MoSeq* pipeline. Analyzing 80,000 frames of open-field or Morris water maze recordings with the traditional pipeline required ~10 hours of computation and ~6 hours of human labor, including 800 annotated frames across 3 anatomical and 8 environmental landmarks over 8 refinement rounds, followed by an additional 4 hours of computation and 0.5 hours of manual effort for behavior classification. In total, this amounted to more than 10 hours of processing with 6.5 hours of human engagement. By contrast, *CASTLE* completed the same dataset in ~1.6 hours with only 15 minutes of user input for ROI annotation and parameter tuning. This dramatic efficiency gain derives from *CASTLE*'s use of pre-trained foundation models, which eliminate the need for extensive manual keypoint labeling and task-specific retraining, thereby demonstrating its scalability for high-throughput behavioral analysis.

### Code availability

The *CASTLE* pipeline and graphical user interface are freely available as open-source software at https://github.com/CASTLE-ai/castle-ai.

The repository provides:

- Complete source code for the *CASTLE* pipeline and GUI
- Installation instructions and dependency specifications
- Links to required pre-trained model checkpoints
- Integration with *CUDA*-accelerated libraries (cuML) for enhanced performance

The software builds upon and acknowledges contributions from

- https://github.com/yoxu515/aot-benchmark
- https://github.com/facebookresearch/segment-anything
- https://github.com/facebookresearch/dinov2 .

For questions, bug reports, and contributions, users are encouraged to utilize the GitHub repository's issue tracking system. All analysis scripts and workflows used to generate the results presented in this manuscript are reproducible using the provided codebase. We used the following packages in our data analysis: av (12.1.0), CUDA toolkit components (cuda-bindings 12.8.0, cuda-python 12.8.0), cuML (25.2.0), deeplabcut (2.3.10), gradio (4.44.0), h5py (3.13.0), huggingface-hub (0.29.1), matplotlib (3.10.0), numpy (2.1.3), opencv-python (4.11.0.86), pandas (2.2.3), pillow (10.4.0), plotly (5.22.0), scikit-learn (1.6.1), scipy (1.15.2), torch (2.6.0), torchaudio (2.6.0), torchvision (0.21.0), umap-learn (0.5.7) and xformers (0.0.29.post3).

### Methods references


52 Obaid, A. et al. Massively parallel microwire arrays integrated with CMOS chips for neural recording. Sci Adv 6, eaay2789 (2020). https://doi.org/10.1126/sciadv.aay2789

53 Pachitariu, M., Steinmetz, N., Kadir, S., Carandini, M. & Kenneth D, H. Kilosort: realtime spike-sorting for extracellular electrophysiology with hundreds of channels. BioRxiv, 061481 (2016).
